# mTORC1 controls phase-separation and the biophysical properties of the cytoplasm by tuning crowding

**DOI:** 10.1101/073866

**Authors:** M. Delarue, G.P. Brittingham, S. Pfeffer, I.V. Surovtsev, S. Ping-lay, K.J. Kennedy, M. Schaffer, J.I. Gutierrez, D. Sang, G. Poterewicz, J.K. Chung, J. Plitzko, J.T. Groves, C. Jacobs-Wagner, B.D. Engel, L.J. Holt

## Abstract

Macromolecular crowding has a profound impact on reaction rates and the physical properties of the cell interior, but the mechanisms that regulate crowding are poorly understood. We developed Genetically Encoded Multimeric nanoparticles (GEMs) to dissect these mechanisms. GEMs are homomultimeric scaffolds fused to a fluorescent protein. GEMs self-assemble into bright, stable fluorescent particles of defined size and shape. By combining tracking of GEMs with genetic and pharmacological approaches, we discovered that the mTORC1 pathway can tune the effective diffusion coefficient of macromolecules ≥15 nm in diameter more than 2-fold without any discernable effect on the motion of molecules ≤5 nm. These mTORC1-dependent changes in crowding and rheology affect phase-separation both *in vitro* and *in vivo.* Together, these results establish a role for mTORC1 in controlling both the biophysical properties of the cytoplasm and the phase-separation of biopolymers.

## Introduction

Molecular crowding is crucial for the efficient function of biological systems (Zhou et al., 2008). If *Xenopus* egg extracts are diluted by more than a few percent, fundamental biological processes such as mitosis and DNA replication fail (Lohka and Maller, 1985). High concentrations of crowding agents entropically favor molecular association events, thereby accelerating molecular reactions (Rivas et al., 2001; Zhou et al., 2008). However, excessive crowding can also dramatically decrease molecular motion, just as the loss of a lane on a freeway can transform smooth traffic flow to instant gridlock (Miermont et al., 2013; Trappe et al., 2001). This kind of jamming depends strongly on particle size: molecules with sizes equivalent to or larger than the dominant crowding agent will be more affected than small particles that can move through the gaps left at the intersections of jammed crowding particles. Thus, changes in molecular crowding can have profound effects on cell physiology and may affect some pathways disproportionately, depending on the sizes of the molecules involved.

A key example where regulation of macromolecular crowding is paramount is phase-separation. Proteins that have a stronger propensity to self-associate than to interact with the solute can undergo a phase transition, where a large number of interacting proteins coalesce into a condensed liquid phase that is separate from the surrounding bulk liquid solute (Banani et al., 2016; Brangwynne et al., 2009). These biological condensates are increasingly observed in diverse fields including cell division (Woodruff et al., 2017; 2015), development (Brangwynne et al., 2009; Smith et al., 2016), cancer (Grabocka and Bar-Sagi, 2016; Kaganovich et al., 2008), neurodegenerative disease (Kwon et al., 2014), T-cell activation (Alberti and Hyman, 2016; Su et al., 2016), and even photosynthesis (Freeman Rosenzweig et al., 2017). Macromolecular crowding tunes phase-separation *in vitro* (Banani et al., 2016). However, the physiological mechanisms that control crowding within the cell and the effects of crowding on cellular processes remain obscure.

One method to study macromolecular crowding and other cellular biophysical properties is to observe the motion of nanoscale tracer particles as they move within the cell. This approach, known as passive microrheology, can be used to infer the viscosity, elasticity, structure, and dynamics of the surrounding material from the characteristic motion of these tracer particles (Mourão et al., 2014; Wirtz, 2009). Various groups have studied the motion of non-biological nanoparticles in cells (Daniels et al., 2006; Luby-Phelps et al., 1986), but these techniques are labor intensive and typically perturb the cell. For example, microinjection disrupts the cell membrane and cortex, and is not feasible in organisms with a cell wall, such as budding yeast. An alternative approach is to track the motion of endogenous structures, such as mRNA molecules tagged with specific loops and loop-binding proteins that can be tagged with fluorescent proteins (Shav-Tal et al., 2004). However, if the motion of an endogenous molecule is affected by a perturbation, it is difficult to know if these changes in motion are due to impacts on the biophysical properties of the cell, or rather caused by direct regulation of the tracer particle. Evolutionarily orthogonal biological tracers have been used to address this issue, notably the μNS particle from mammalian rheovirus (Joyner et al., 2016a; Parry et al., 2014). These types of tracer particles are less likely to undergo specific regulated interactions with the cell, but a major limitation of the μNS system is that these condensates do not have a predefined size, and thus require additional calibration steps to convert fluorescence measurements into particle size (Parry et al., 2014). Furthermore, the size of μNS probes (> 50 nm) is larger than most multimeric protein complexes found inside cells.

In order to address these issues, we developed Genetically Encoded Multimeric (GEM) nanoparticles (henceforth GEMs), which are bright tracer particles of a defined shape and size. GEMs can serve as a standard microrheological tool across a broad range of organisms; in this study, we used GEMs in *S. cerevisiae* and human cell lines. By using GEMs from a different kingdom than the organism of study, we make it far less likely that the particles will be affected by specific interactions. With this technology in hand, we screened for mechanisms that regulate the biophysical properties of cells. We found that mTORC1 controls ribosome abundance through a combination of cell volume control, ribosome biogenesis and autophagy. *In situ* cryo-electron tomography (cryo-ET) of the native cellular environment (Asano et al., 2016) revealed that inhibition of mTORC1 nearly halves the cytosolic ribosome concentration. As ribosomes account for ~20% of the total cytosolic volume, modulation of their concentration has a dramatic effect on the biophysical properties of the cell. This modulation is significant: Inhibition of mTORC1 can double the effective diffusion coefficient of particles that are 15 nm in diameter or greater. We derived a theoretical model based on the phenomenological Doolittle equation that relates the diffusion of a tracer particle to the fraction of crowding close to a jamming transition, and were able to predict changes in the diffusion coefficient as a function of ribosome concentration in both budding yeast (S. *cerevisiae)* and human cells (HEK293). Finally, we found that changes in macromolecular crowding downstream of mTORC1 tune phase-separation in both yeast and human cells, providing a direct link between *in vivo* crowding regulation and phase-separation.

## Results

### GEMs can be made from both 15 nm and 41 nm icosahedral protein cages

We developed GEMs to study the rheological properties of the eukaryotic cytoplasm. We began with natural homomultimeric scaffolds that self-assemble into icosahedral geometries and fused these scaffolds to fluorescent proteins to create fluorescent GEMs.

In this study, we employed two GEMs of 40 nm and 15 nm diameter, with scaffolding domains based on the encapsulin protein from the hyperthermophilic archaeon *Pyrococcus furiosus* (Akita et al., 2007) and the lumazine synthase enzyme complex from the hyperthermophilic bacterium *Aquifex aeolicus* (Zhang et al., 2001), respectively (figure 1A-C). When expressed within cells, these GEMs self-assembled into bright, stable particles (figure 2A-B).

**Figure 1.**
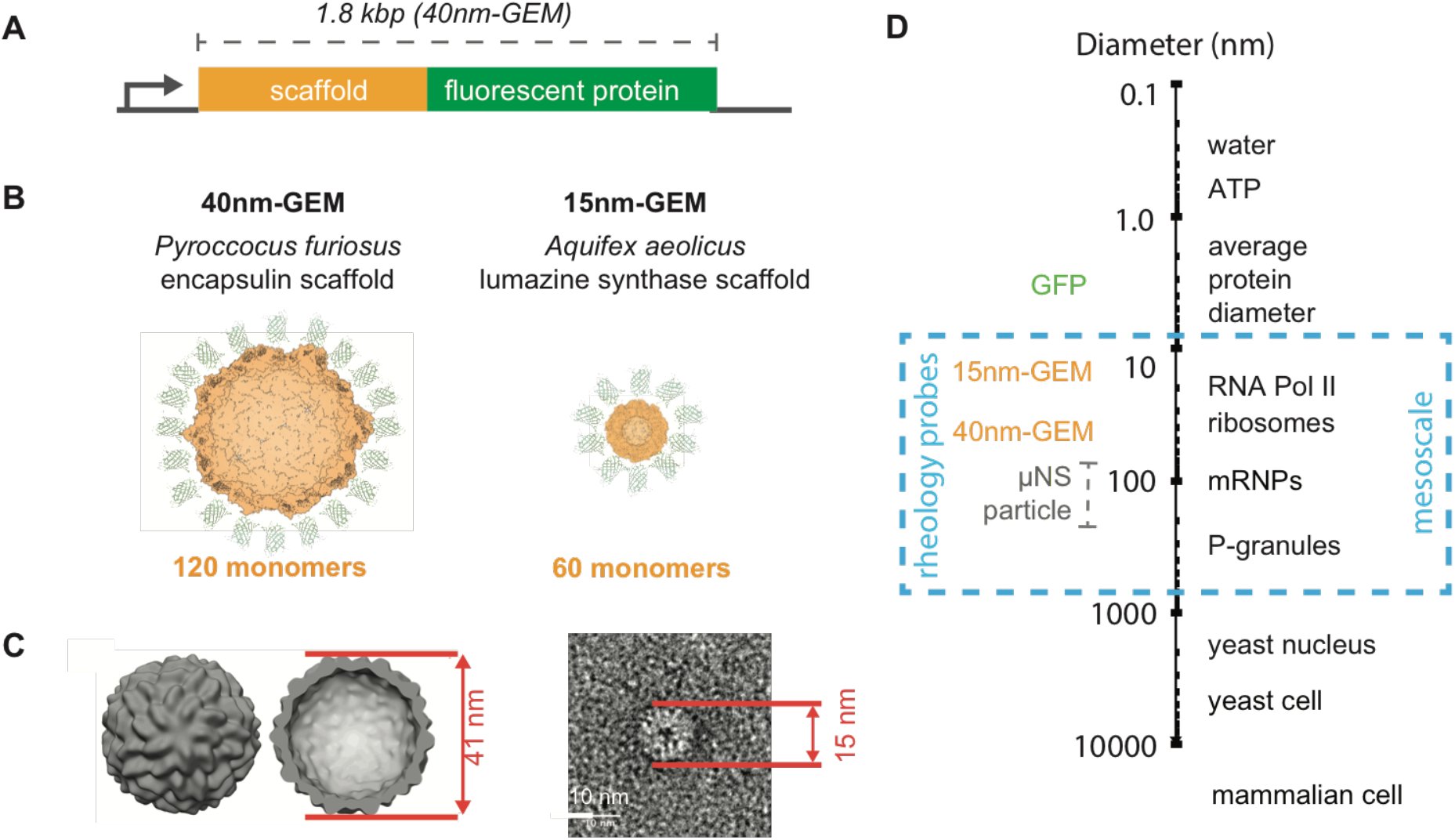
Genetically Encoded Multimeric nanoparticles (GEMs) are homomultimeric fluorescent nanoparticles that self-assemble to a stereotyped size and shape. **(A)** General gene structure of GEMs, which consist of an in-frame fusion of a multimerizing scaffold to a fluorescent protein. **(B)** Predicted structures of 40nm-GEMs and 15nm-GEMs. **(C)** Left, cryo-ET subtomogram average of 40nm-GEMs within the cell; Right, averaged negative stain EM images of 15nm-GEMs. **(D)** Diameters of GEMs and other macromolecules at the meso length-scale, shown in relation to small molecules, protein complexes, and cells.

**Figure 2.**
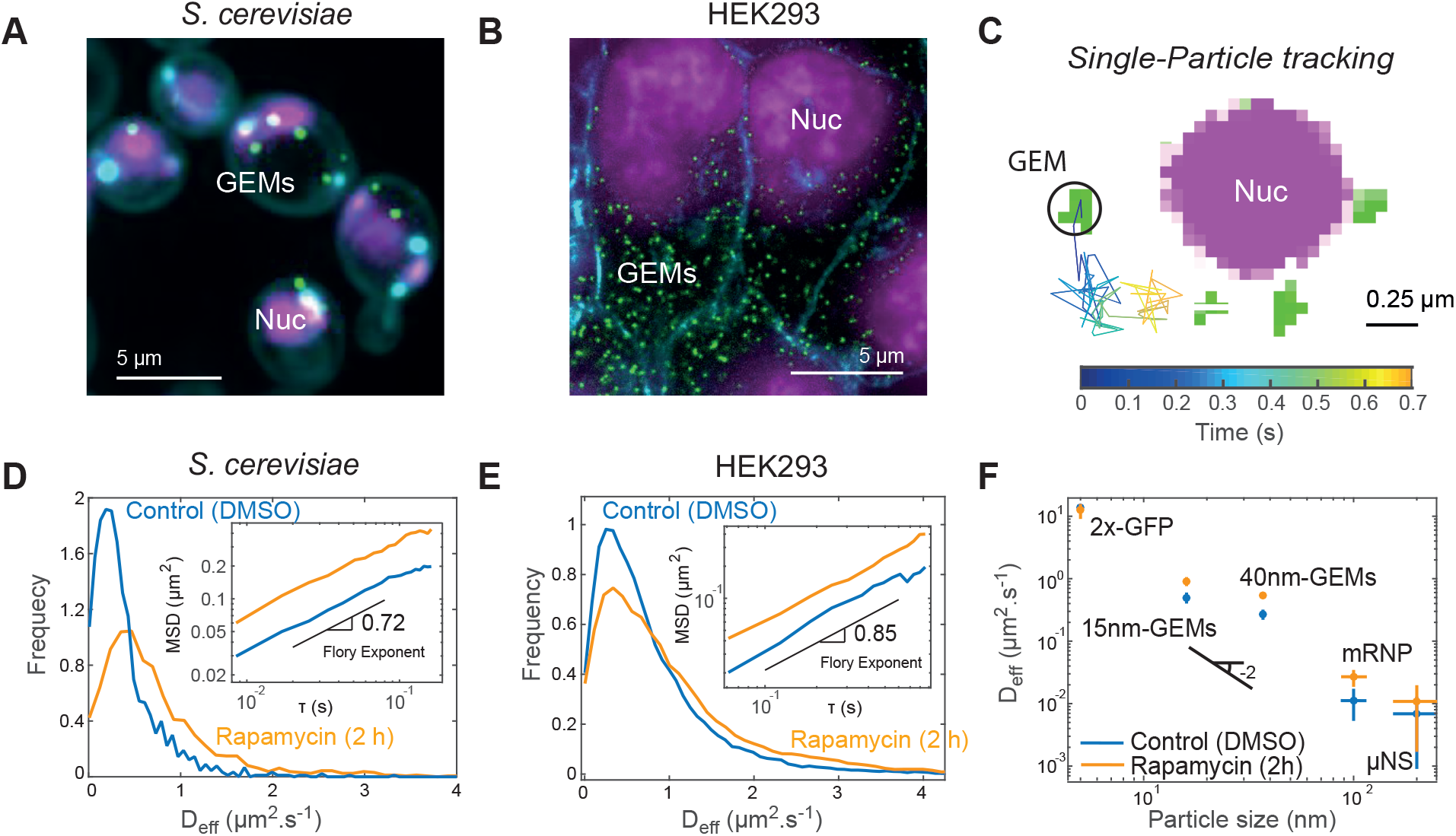
mTORCI inhibition increases the effective diffusion coefficient of GEMs. **(A)** 40nm-GEMs expressed in **(A)** *S. cerevisiae* and **(B)** HEK293 cells. GEMs are visualized using a T-Sapphire fluorescent protein (green). DNA is visualized with SiR-Hoeschst (magenta). Yeast cell walls and mitochondria were visualized using calcoflu-or-white and HEK293 membrane with wheat germ agglutinin (cyan). **(C)** High magnification example of tracking of a 40-nm GEM particle (green) within an *S. cerevisiae* cell, imaging at 100 frames per second. Three other GEMs and the nucleus (magenta) are also displayed. (D-E) Distribution of 40nm-GEM effective diffusion coefficients (D_eff_) for *S. cerevisiae* (**D**) and HEK293 (**E**); results from DMSO (carrier control) treatment are displayed in blue; rapamycin treatment results are displayed in orange. Insets: time and ensemble-averaged mean-square displacements in log-log space with the Flory exponent indicated. (**F**) Effect of rapamycin on the effective diffusion coefficients of endoge-nous molecules and tracer particles of various sizes. Indicated, the -2 power-law scaling of diffusion coefficient as a function of diameter, which does not conform to Stokes-Einstein predictions.

Using *in situ* cryo-ET to image the native cellular environment (Asano et al., 2016), we determined that the *Pyrococcus furiosus* encapuslin GEMs formed assemblies with a diameter of 41 nm, a little larger than the 35 nm diameter reported from crystallography data (Akita et al., 2007) (figure 1C). This larger diameter is likely due to the additional GFP molecules decorating the encapsulin particle. We noted that there was no size variation within cells of 40 nm GEMs in any condition that we inspected (figure S3). Thus, we termed these particles **40nm-GEMs**.

Using negative stain electron microscopy, we determined that the *A. aeolicus* lumazine synthase protein assembles into 15 nm particles (figure 1C), in good agree ment with crystallography data (Zhang et al., 2001) (figure 1C). Thus, we termed these particles **15nm-GEMs**.

The 15nm-GEMs and 40nm-GEMs are in the range of multi-subunit assemblies such as ribosomes, proteasomes and chromatin remodeling complexes (figure 1D), thus allowing us to investigate the mesoscale microrheological environment experienced by these complexes. Therefore, these biologically-orthogonal nanoparticles probe the biophysical properties of the cell at a length-scale that was previously inaccessible.

### GEMs allow rapid characterization of the rheological properties of the cytosol in yeast and human cells

We expressed 40nm-GEMs in the budding yeast *S. cerevisiae* and an adenovirus transformed Human Embryonic Kidney cell line (HEK293, (Russell et al., 1977)) (figure 2A-B, supplemental movies 1-2). 40nm-GEMs are bright enough to allow single particle tracking at 10 ms frame rates (figure 2C, see methods). We compared thousands of individual traces to extract the effective coefficient of diffusion, D_eff_, at short timescales. GEM motilities differ between the two biological systems: 40nm-GEMs have a median effective diffusion coefficient of ~0.3 μm^2^ s^-1^ in yeast and ~0.5 μm^2^ s^-1^ in mammalian cells (figure 2D-E). These estimates are in good agreement with expectations from the literature (Luby-Phelps et al., 1986), further supporting their use as micro-rheological standards. Using time and ensemble-averaging, we inspected the mean-squared displacement (MSD) curves at longer timescales and found that 40nm-GEMs were subdiffusive (inset figure 2D-E) with a Flory exponent of ~0.72 in yeast and ~0.85 in HEK293 cells. This subdiffusive motion could be due to local caging within a highly crowded environment close to a glass transition and/or interactions between the tracer particle and the environment (Wang et al., 2012). However, the Flory exponent did not change significantly in most of our perturbation experiments, so we focused on the apparent diffusion coefficient as our main metric to report on cytosolic rheology.

### mTORC1 affects the biophysical properties of the cytosol

In initial experiments in yeast, we observed that cell culture conditions changed the apparent diffusion coefficients of 40nm-GEMs. When yeast cultures approached saturation, the effective diffusion rates of GEMs increased (not shown). By supplementing and depleting various growth medium components, we found that this increase in effective diffusion occurred in response to amino acid depletion (not shown).

The mechanistic target of rapamycin complex (mTORC1) is the major amino acid sensor in *eukaryotes* (Hara et al., 1998). Therefore, we hypothesized that mTORC1 signaling might cause the observed changes in cytoplasmic rheology in response to perturbations in amino acid levels. mTORC1 can be inhibited by addition of rapamycin, which forms an inhibitory complex with the protein FKBP12 (encoded by *FPR1* in *S. cerevisiae)* (Heitman et al., 1991). Consistent with our hypothesis, 40nm-GEMs displayed increased mobility upon inhibition of mTORC1 with rapamycin in both *S. cerevisiae* and HEK293 cells (figure 2D-E, supplemental movies 1-2). Changes in the distribution of diffusion coefficients were highly significant (p < 1 x 10^−41^ for yeast and p < 1 x 10^−40^ for HEK293; Kolmogorov-Smirnov test). These results suggest that mTORC1 controls the biophysical properties of the cytosol at the 40 nm length-scale in both yeast and mammalian cells.

### mTORC1 does not affect diffusion at the length-scale of individual proteins

The change in effective diffusion of 40nm-GEMs was abundantly clear, but rheology can vary considerably between different length-scales. Therefore, we studied other particles to check the generality and length-scale dependence of the changes in microrheology downstream of mTORC1 signaling. First, we repeated our experiments with 15nm-GEMs and found that their diffusion also increased upon mTORC1 inhibition (figure 2F). We also saw an increase in the diffusion coefficients of larger structures, such as the endogenous *GFA1* mRNP tagged with the PP7-GFP system (Joyner et al., 2016b) and GFP-μNS particles (figure 2F). These particles are approximately 100 nm and 200 nm in diameter, respectively. Thus, mTORC1 modulates the effective diffusion coefficient of particles in the mesoscale 15 to 200 nm diameter range.

To probe rheology at shorter length-scales, we used fluorescence correlation spectroscopy to calculate the effective diffusion of a double-GFP molecule, which is around 5 nm in diameter. The diffusion of this smaller protein was unaffected by the addition of rapamycin (figure 2F, S7, Table S1). Thus, mTORC1 inhibition increases the diffusion coefficients of particles at or above the typical size of multimeric protein complexes, but particles that are the typical size of monomeric proteins or smaller are unaffected.

### Changes in cell size, translation or the cytoskeleton do not account for the effects of mTORC1 on the motion of 40nm-GEMs

Rapamycin treatment arrests cell division, but does not completely prevent cell growth. Thus, cell volume continues to increase (Chan and Marshall, 2014). Therefore, we hypothesized that the increase in the effective diffusion coefficients of 40nm-GEMs might be through increases in cell volume. To test this idea, we took advantage of a chemical genetic strategy that involves inhibition of a *cdc28-as1* allele of budding yeast Cyclin Dependent Kinase 1 (Cdk1) with 1-NM-PP1 (Bishop et al., 2000). Upon complete inhibition of *Cdk1-as1* with 10 μM 1-NM-PP1, cell division arrested and cytoplasmic volume increased, but no changes were observed in the motion of 40nm-GEMs (Figure S1A-B). Thus, cell volume increase is not sufficient for the observed biophysical effects.

Protein translation is regulated by mTORC1: when nutrients and growth factors are present, cells enter an anabolic state and protein translation is upregulated in an mTORC1-dependent manner. Inhibition of mTORC1 with rapamycin leads to rapid inhibition of translation. Therefore, we tested whether decreases in translation could explain the observed changes in the effective diffusion coefficients of 40nm-GEMs. To investigate this idea, we stalled translation by addition of 1 μM cycloheximide. The median half-life of approximately 4,000 yeast proteins is about 40 minutes under these conditions (Belle et al., 2006). The motion of 40nm-GEMs was neither affected during acute cycloheximide treatment, nor after 180 minutes of treatment (Figure S1A-B). These results suggest that neither translational inhibition nor protein degradation explain our observations.

Another plausible hypothesis was that mTORC1 might alter the dynamics or structure of the cytoskeleton. We used Latrunculin A to depolymerize the actin cytoskeleton (figure S1C) and found that, while the basal rate of 40nm-GEM diffusion did decrease, there was no epistasis with the rapamycin effect. This result suggests that the actin cytoskeleton contributes to the fluidity of the *S. cerevisiae* cytoplasm, but that mTORC1 does not modulate mesoscale rheology through actin-dependent effects. We then used nocodazole to depolymerize microtubules and found that the basal rate of 40nm-GEMs slightly increased. Here there was some epistasis with rapamycin, but the effect size was small compared to other mutants (see below). Thus, the actin and microtubule cytoskeleton play an important role in defining the mesoscale properties of the yeast cytoplasm, but do not appear to be the primary mechanistic explanation for the regulation of rheology by mTORC1.

### mTORC1 controls cytoplasmic rheology by tuning ribosome concentration

In our experiments in *S. cerevisiae*, we typically saw >10 40nm-GEMs in each cell, and imaged at ~100 Hz. In this manner, we collected thousands of traces in a matter of seconds. Because every cell expressed GEMs, there was no time delay associated with finding cells, and no laborious and disruptive manipulations like microinjection. These advantages enabled us to use GEMs for high-throughput screens to determine the mechanisms that control the biophysical properties of the cell.

We screened >100 candidate mutants in *S. cerevisiae* to investigate how mTORC1 might control cellular rheology (selected genes are listed in Table S2). We used an *fpr1Δ* strain as a negative control. *FPR1* encodes the immunophilin protein FKBP12 that binds to rapamycin. It is the FKBP12-rapamycin complex that binds and inhibits the mTORC1 complex (Heitman et al., 1991). Thus, in the absence of the *FPR1* gene, rapamycin cannot inhibit mTORC1. In accordance with this expectation, there was no detectable effect of rapamycin on the *fpr1Δ* deletion strain (figure 3A).

**Figure 3.**
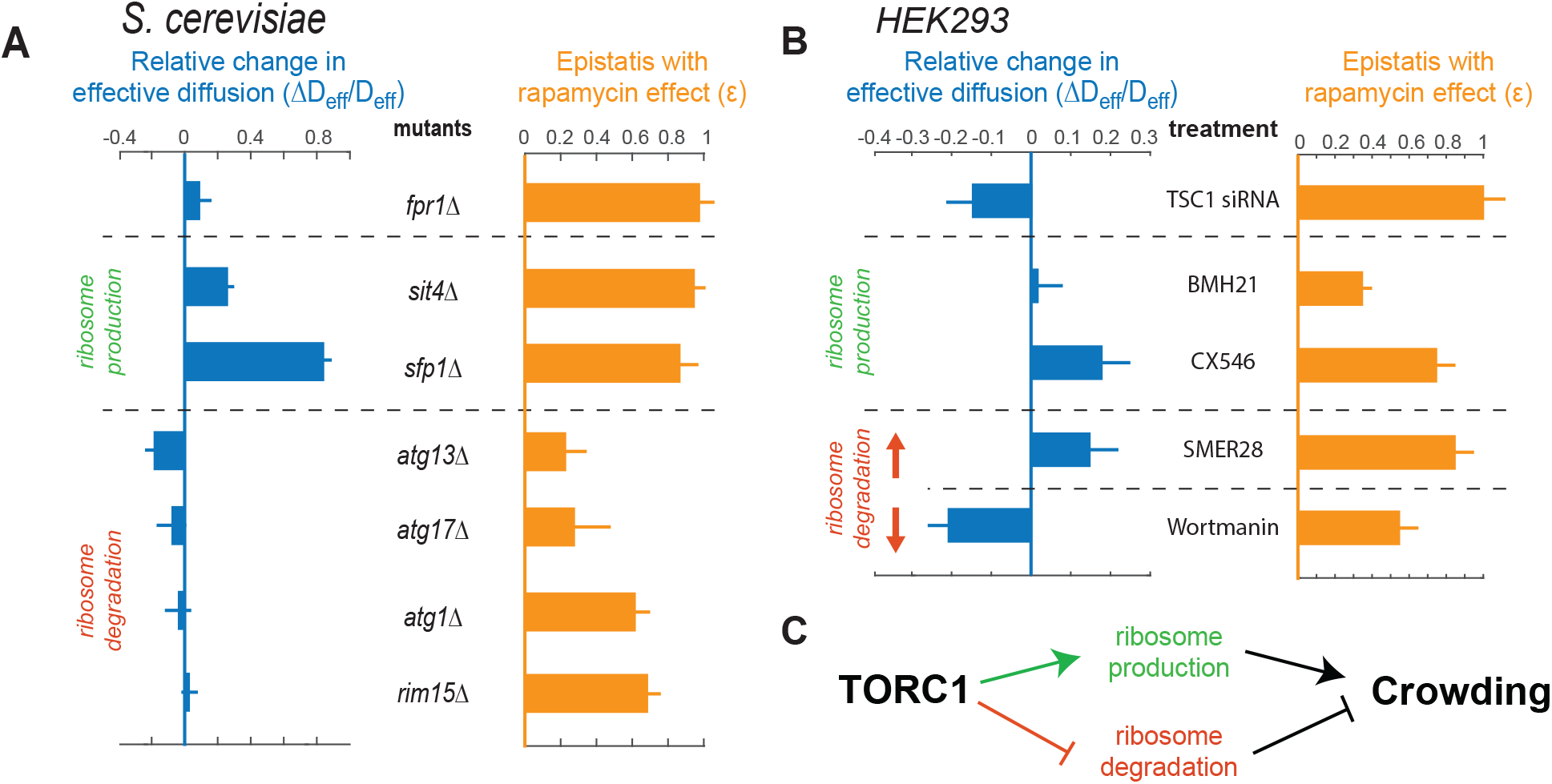
mTORC1 controls the effective diffusion coefficient of 40nm-GEMs by tuning ribosome concentration. (**A**) Selected mutants from a candidate screen in *S. cerevisiae.* The change in the baseline effective diffusion coefficients of 40nm-GEMs (left, blue) is plotted for each mutant; along with the epistasis between these conditions and rapamycin treatment (right, orange). (**B**) Pharmacological and siRNA perturbations in HEK293 cells suggest that mTORC1 also modulates cytoplasmic rheology through ribosome crowding in mammals. Data are presented as the median +/-SEM (standard error of the mean). (**C**) Proposed model of crowding control in *S. cerevisiae* and HEK293 cells.

The *SIT4* gene encodes a subunit of the PP2A phosphatase required for a major signaling branch downstream of mTORC1 (Di Como and Arndt, 1996; Peterson et al., 1999a). The *sit4Δ* mutation was epistatic to the rapamycin treatment, as addition of rapamycin to *sit4Δ* cells had little to no effect on particle diffusion. Together, these results validated the use of 40nm-GEMs in genetic screens and constrained our genetic screen to the PP2A-dependent branch of mTORC1-signaling.

We tested and rejected several hypotheses for the possible mechanism through which mTORC1 signaling might affect cytosolic biophysics (table S2). Eventually, we found that deletion of the *SFP1* gene, which encodes a transcription factor involved in ribosomal RNA biogenesis (Fingerman et al., 2003) affected the effective diffusion coefficient of 40nm-GEMs even more than rapamycin treatment (figure 3A, left). Furthermore, the *sfplΔ* deletion strain had complete epistasis with rapamycin (figure 3A, right). These results implicated ribosome biogenesis as a key mechanism in the control of cellular rheology.

The steady-state concentration of ribosomes in the cytoplasm is determined by the rate of ribosomal production, which is strongly affected by the *SIT4* and *SFP1* genes (Peterson et al., 1999b), and the rate of ribosomal degradation. Ribosomes are usually quite stable, but starvation conditions can drive autophagy and ribophagy to accelerate rates of ribosome degradation, especially when mTORC1 is inhibited (Waliullah et al., 2017). This starvation response is thought to scavenge cellular macromolecules and organelles to recycle cellular building blocks (Reggiori and Klionsky, 2013), but reduction in the concentration of ribosomes has also been proposed as a function for these pathways (Tsukada and Ohsumi, 1993). In accordance with this latter idea, mutations in the autophagy genes *ATG1, ATG13 and ATG17* and the ribophagy gene *RIM15* (Waliullah et al., 2017) all showed significant epistasis with rapamycin treatment (figure 3A, right).

Next, we sought to determine whether the mechanisms that we identified in *S. cerevisiae* would also hold true in mammalian cells. To this end, we employed HEK293 cells stably transduced or transfected with 40nm-GEMs and used pharmacological perturbations and siRNA to test whether ribosome concentration was important in setting the biophysical properties of mammalian cells at the 40 nm length-scale.

Inhibition of ribosome production using the small molecules BMH-21 (Peltonen et al., 2010) or CX5461 (Drygin et al., 2011) showed strong epistasis with the effect of rapamycin (figure 3B, right). However, the basal diffusion coefficient only increased in CX5461 treatment (figure 3B, left). We speculate that the failure of BMH-21 to impact GEM motion could be due to the pleiotropic nature of this drug, which could cause compensatory effects in the basal biophysical properties of the cytoplasm. Nevertheless, these pharmacological perturbations suggest that control of rRNA transcription is part of the mechanism by which mTORC1 inhibition increases the fluidity of mammalian cells.

Stimulation of autophagy using the SMER28 compound (Tian et al., 2011), thereby reducing ribosome concentration, led to an increase in the basal diffusion of 40nm-GEMs (figure 3B, left) and exhibited strong epistasis with rapamycin (figure 3B, right). In contrast, decreasing autophagy with Wortmanin, which is predicted to increase ribosome concentration (Hansen et al., 1995), led to decreased basal diffusion (figure 3B, left) and epistasis with rapamycin (figure 3B, right).

Finally, we increased mTORC1 activity by siRNA-mediated knock-down of the mTORC1 inhibitor TSC1 (Potter et al., 2001). This treatment led to a decrease in basal diffusion (figure 3B, left, S8D). Thus, after screening over 100 mutants and drug treatments, we found that the conditions that most strongly affected the baseline rate of GEM diffusion and/or showed epistasis with rapamycin treatment fell into two general classes: ribosome biogenesis and autophagy.

Together, these data suggested that mTORC1 controls macromolecular crowding by tuning ribosome concentration (figure 3C).

### Ribosomes act as crowding agents

Ribosomes are one of the most abundant macromolecules in the cytoplasm (Duncan and Hershey, 1983; Warner, 1999). Our genetic and pharmacological results suggested that ribosomes are the main crowding agent regulated by mTORC1. To test this hypothesis, we counted ribosomes within the native cellular environment with single molecule precision.

We used *in situ* cryo-ET to directly visualize ribosomes within the yeast cytoplasm in control and rapamycin-treated conditions. Briefly, we thinned vitreous frozen yeast cells by focused ion beam (FIB) milling (Marko et al., 2007; Rigort et al., 2012; Schaffer et al., 2017) and then performed *in situ* cryo-ET (Asano et al., 2016) to produce three-dimensional images of the native cellular environment at molecular resolution (figures 4A-B, S2–S6, supplemental movies 3-4). Template matching enabled us identify ribosomes within the cellular volumes with high sensitivity (figure S2). Subsequent subtomogram averaging produced *in situ* structures of the ~30 nm ribosomes and 40nm-GEMs at 11.5 Å and 26.3 Å resolutions, respectively (figures 4C, S2–S6). In W303 yeast cells undergoing log phase growth, the concentration of ribosomes in the cytoplasm was ~14,000 ribosomes/μm^3^ (23 μM), whereas this concentration decreased almost 2-fold to ~8,000 ribosomes/μm^3^ (13 μM) when cells were treated with rapamycin for two hours (figure 4D). This corresponds to a drop from ribosomes occupying ~20% of the cytosolic volume to ~12%.

**Figure 4.**
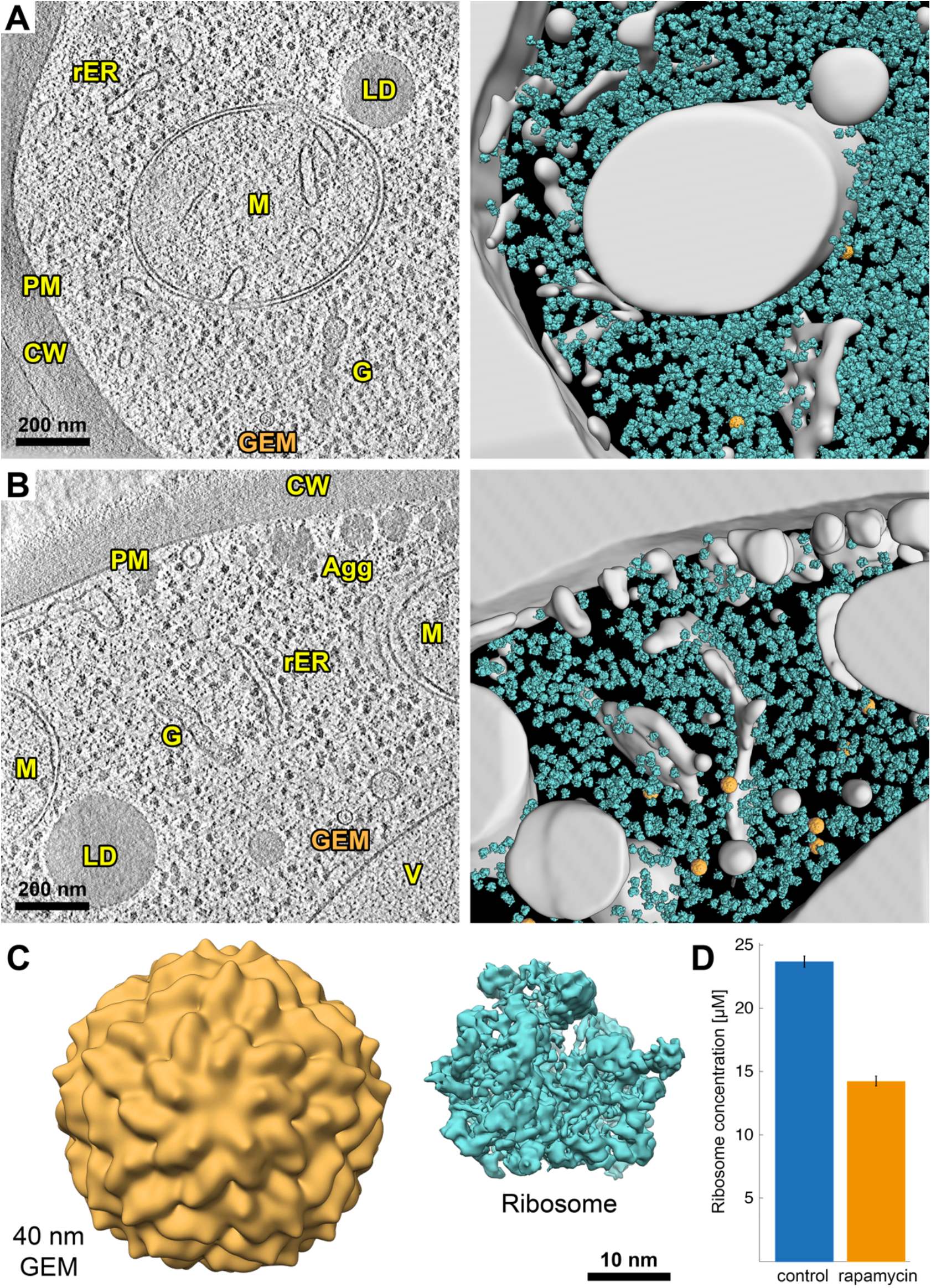
*In situ* cryo-electron tomography of FIB-milled *S. cerevisiae* reveals that ribosome concentration dramatically decreases upon mTORC1 inhibition. (**A**) DMSO-treated cells. **(B)** Rapamycin-treated cells. (**Left**) Slice through a representative cryo-electron tomogram of a FIB-milled yeast cell. The cell wall (CW), plasma membrane (PM), rough endoplasmic reticulum (rER), lipid droplets (LD), mitochondria (M), Golgi apparatus (G), vacuole (V), aggregates (Agg), and one example GEM nanoparticle are indicated. (**Right**) 3D segmentation of the same tomogram showing ribosomes (cyan) and GEMs (orange). The non-cytosolic volume is grey. **(C)** Subtomogram averages of the 40nm-GEM nanoparticles and ~30 nm ribosomes from within the cellular volumes, shown in relative proportion. **(D)** Cytosolic ribosome concentrations after 2 h DMSO (blue) and rapamycin (orange) treatment. Error bars are SEM. Concentrations were calculated from 14 DMSO-treated and 13 rapamycin-treated cells (see Figs. S5 and S6).

### Ribosomes control the biophysical properties of the cytosol

Together, the screens and *in situ* cryo-ET data together strongly suggested a causal relationship between ribosome concentration and the motion of particles at the 15 nm and 40 nm length-scales. We therefore developed a physical model based on the phenomenological Doolittle equation to predict the effective diffusion coefficients of particles as a function of ribosome concentration (see methods, equation S5. Briefly, the model predicts the effective diffusion coefficient of a particle as a function of the volume fraction of crowders inside the cell (figure 5A). A key assumption of the Doolittle equation as that the volume fraction of crowder is close to a jamming transition, such that a small increase of the concentration of crowding agent leads to a dramatic decrease in the effective diffusion coefficient. This model is constrained by two parameters: the non-osmotic volume *v*\*, which is the volume occupied by all macromolecules when no further water can be osmotically extracted from the cell, and a prefactor parameter denoted *ζ* which most likely relates to the degree of interaction of the tracer particle with its surrounding microenvironment.

**Figure 5.**
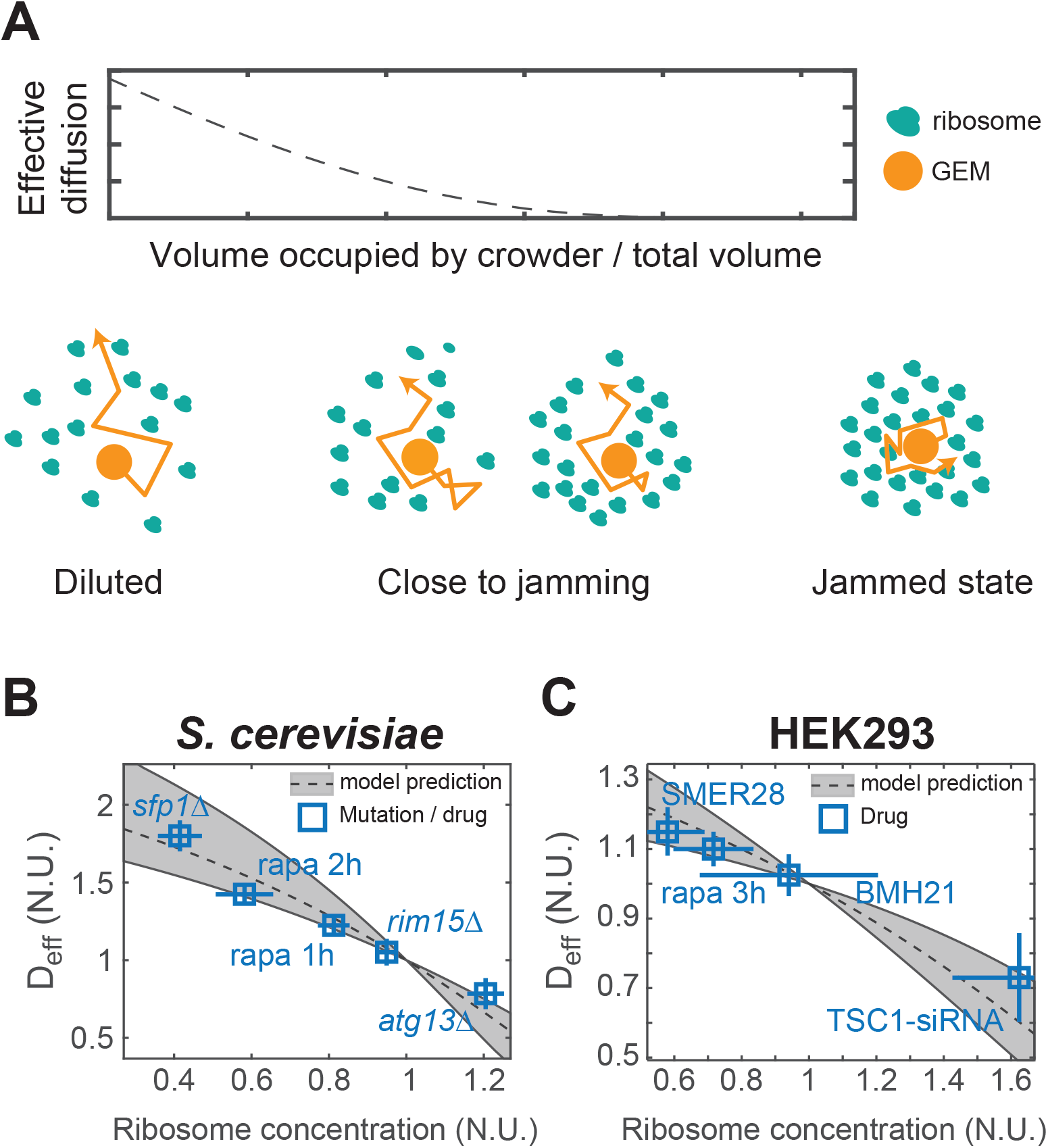
A physical model of the cytosol near to a jamming transition accurately predicts diffusion as a function of ribosome concentration. (**A**) The phenomenological Doolittle equation describes the effective diffusion coefficient of particles as a function of excluded volume when the system is close to the jamming transition. (**B** and **C**) A model based on the Doolittle equation, and parameterized empirically with no fitted parameters, accurately predicts the diffusion coefficient of 40nm-GEMs in both yeast (**B**) and HEK293 cells (**C**) as a function of the concentration of ribosomes (measured by quantification of a total extracted nucleic acids, see Figure S8E-G.) Median coefficients of diffusion are normalized to WT conditions on the day the data was acquired and plotted as the median +/-SEM.

We empirically determined both of these parameters (*v*\* and *ζ*) using instantaneous cell volume changes through osmotic perturbations (figure S8A-B). We found that the non-osmotic parameter is smaller for HEK293 than for *S. cerevisiae*, confirming our expectation that HEK293 cells are less crowded than yeast. The parameter *ζ* is very similar in both species, perhaps suggesting that 40nm-GEMs have similar interactions with the human and yeast cytosol, a result most easily explained by GEMs having very little specific interaction with their local environment. This concordance further supports the use of GEMs as a microrheological standard across organisms.

Once we had determined the parameters *v*\* and *ζ*, we were able to predict the effective diffusion coefficient of GEMs as a function of ribosome concentration (figure S8E-G; see methods, equation S12). We plotted this prediction for both budding yeast and mammalian cells (figure 5B-C) and found excellent agreement between our predictions and data from control conditions, rapamycin conditions and various genetic and chemical perturbations. In all cases, our model was able to accurately predict the coefficient of diffusion for GEMs over a wide range of ribosome concentrations. All parameters were experimentally determined and no data fitting was required.

We also experimentally determined the prefactor *ζ* for the endogenous *GFA1* messenger ribonucleoprotein complex (mRNP) tagged with the PP7-GFP system. These particles are approximately 100-200 nm in diameter. Once we determined *ζ* for *GFA1* mRNP particles, our model accurately predicted their effective diffusion coefficient as a function of ribosome concentration (figure S8C). Therefore, our results strongly suggest that macromolecular crowding by ribosomes described as hard spheres plays a significant role in determining the biophysical properties of the cytosol at the 40 to 200 nm length-scale.

### mTORC1 tunes phase-separation by controlling ribosome concentration

Macromolecular crowding favors the interaction of macromolecules through depletion attraction effects (Mourão et al., 2014). When large numbers of multivalent proteins exceed a critical nucleation concentration they can condense to form a phase-separated liquid droplet. These liquid droplets can further mature to form gels and amorphous aggregates, including pathogenic amyloid fibers and prions (Alberti and Hyman, 2016).

The phase-separation of biomolecules is tuned by multiple physicochemical effects including the association and dissociation constants of interaction domains, the strength of the interaction of each molecule with the solute phase, and attraction depletion effects that can entropically favor condensation. This latter effect is strongly influenced by macromolecular crowding. Since ribosomes are the dominant crowding agent in the cytoplasm, we hypothesized that ribosome concentration tunes phase-separation. To test this idea, we took advantage of a synthetic system that is well characterized in terms of physicochemical parameters and that phase-separates into liquid droplets both *in vitro* and *in vivo.* This system is comprised of ten repeats of the small ubiquitin-like modifier (SUMO) domain and six repeats of sumo interaction motif (SIM), and has been proven as a reliable model for phase-separation (Banani et al., 2016).

We assessed the effects of ribosomes on the phase-separation of a decamer of SUMO (SUMO10), which can condense with a hexamer of SIM (SIM_6_). Beginning *in vitro*, we added ribosomes purified from *Escherichia coli* over a biologically-relevant concentration range determined from our cryo-ET experiments. We observed that the concentration of SUMO10 and SIM_6_ that partitioned into the condensed liquid droplet phase (partition coefficient) increased as ribosome concentrations increased. Indeed, the partition coefficient increased >50% when ribosome concentration was increased from 13 μM (mTORC1 inhibition) to 23 μM (log-phase growth) (figure 6A).

**Figure 6.**
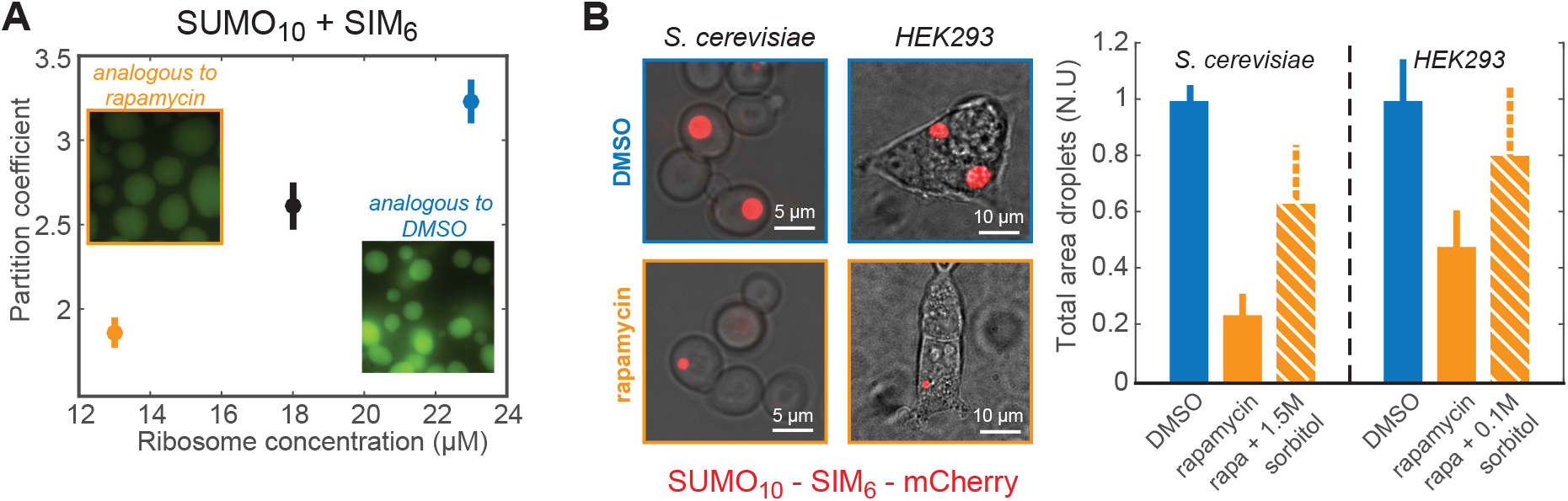
Ribosomes act as a crowding agent that drives phase-separation both *in vitro* and *in vivo.* (**A**) A homodecamer repeat of SUMO (SUMO_10_) was mixed with a homohexamer repeat Sumo Interaction Motif peptide (**SIM**_6_) to achieve equimolar concentrations of each monomer (60 *μ*M). SUMO_10_ + SIM_6_ was kept at constant concentration and incubated with an increasing concentration of fully assembled 70S ribosomes (purified from *E. coli).* There was a >50% increase in the partition coefficient of SUMO_10_ + SIM_6_ when ribosome concentration was increased from 13 *μ*M (equivalent to yeast treated with rapamycin) to 23 *μ*M (the concentration of ribosomes in logarithmically growing yeast cells). (**B**) An in-frame fusion of SUMO_10_-SIM_6_-GFP was expressed in budding yeast *(S. cerevisiae* W303) and HEK293 cells. Left: Micrographs of control cells (DMSO) and cells treated with rapamycin for 2 h. Right: Quantification of total area of phase-separated droplets in control cells (blue), cells treated with rapamycin (orange), and cells treated with rapamycin followed with a hyperosmotic shock with 1.5M (yeast cells) or 0.1M (human cells) sorbitol (orange bars with white cross hatches).

Next, we expressed an in-frame fusion of SUMO10 and SIM_6_ (SUMO_10_-SIM_6_) in yeast and HEK293 cells to study the effects of macromolecular crowding on phase-separation *in vivo.* Due to the challenge of defining a partition-coefficient *in vivo*, we measured the total droplet area per cell as a metric of phase-separation. Inhibition of mTORC1 for two hours led to an 80% and 50% decrease in SUMO_10_-SIM_6_ droplet area in yeast and human HEK293 cells, respectively (figure 6B). We were able to partially recover phase-separation in rapamycin-treated cells by using an acute osmotic shock that reduced cell volume to an extent that restored ribosome concentrations to control levels (figure 6B, right; orange cross-hatched bars). This latter result suggests that mTORC1 controls SUMOi_0_-SIM_6_ phase-separation through ribosome concentration and additional mechanisms, perhaps including translation of SUMO_10_-SIM_6_. It is also likely that phase-separation of SUMO_10_-SIM_6_ cannot reach steady-state before the cell begins to adapt to rapamycin and osmotic shock. Thus, the results are consistent with our model but difficult to interpret at a detailed physical level.

Taken together, these data demonstrate that ribosomes act as macromolecular crowders that tune phase-separation. mTORC1 controls ribosome concentration and therefore is predicted to influence the phase-separation of all cytosolic biomolecules through physicochemical effects including depletion attraction and the tuning of cell rheology.

## Discussion

In this study, we discovered that mTORC1 tunes cytoplasmic rheology through control of ribosome concentration. Recent work has reported dramatic changes in cytoplasmic rheology in response to changes in cellular energy state and metabolism. For example, depletion of ATP in *E. coli* leads to a glass-transition that greatly reduces macromolecular mobility (Parry et al., 2014); glucose starvation in yeast leads to a loss of cell volume and a decrease in the effective diffusion of mRNPs and chromosomal loci due to increased molecular crowding (Joyner et al., 2016a); and decreases in cytoplasmic pH lead to a gel transition in the cytosol of yeast, associated with entry into a dormant state (Munder et al., 2016). All of these responses decrease the fluidity of the cytosol. Interestingly, in contrast to carbon starvation, ATP depletion and acidification, we show that inhibition of mTORC1 increases the fluidity of the cytosol.

Using GEM nanoparticles, we were able to determine the mechanism for this biophysical change: ribosome concentration dominates the rheological properties of the cytoplasm at 15 to 200 nm length-scales. Length-scale considerations in cytoplasmic fluidity have interesting implications for previous findings; for example, solidification of the yeast cytoplasm under glucose starvation was observed using tracking of GFP-μNS particles, which are very large condensates. However, it would be surprising if the diffusion of *all* macromolecules decreases. Our results are more consistent with the idea that smaller individual proteins (< 15 nm diameter) can keep moving through the interstitial space between the larger jammed particles (ribosomes). This result is in agreement with the particle size dependency observed in the bacterial cytoplasm (Parry et al, 2014). In a scenario where larger macromolecules are spatially confined while smaller macromolecules continue to diffuse, processes dependent on large complexes, such as apoptosis and translation, may be affected, while many other basic cellular functions can continue unchanged.

Ribosomes are one of the most abundant macromolecules in the cell (around 200,000 ribosomes per yeast cell (Warner, 1999) and 3,000,000 per HeLa cell (Duncan and Hershey, 1983), and we determined that ribosomes occupy 20% of the volume of the cytosol. Indeed, when we model the cytoplasm as a solution filled with ribosomes (assumed to be hard spheres), we can accurately predict the diffusion coefficient of 40nm-GEMs tracer particles and endogenous mRNPs as a function of ribosome concentration. Our model characterizes the cytoplasm close to a jamming transition, which implies that a small variation in water content will have a profound effect on the basal diffusion of protein complexes. A glass transition is one type of jamming transition, therefore our model may help explain some of the observations of glass transitions in previous studies (Joyner et al., 2016a; Parry et al., 2014).

Physiological regulation of the thousands to tens of thousands of different proteins found within cells is a complex task. This regulation is achieved through finegrained mechanisms, including transcriptional and translational control of protein abundance as well as post-translational modifications such as protein phosphorylation and ubiquitylation. However, our studies suggest that macromolecular crowding could also lead to broad regulation of cell state. Changes in macromolecular crowding may provide coarse-grained regulation of protein interactions, diffusion and folding; the cell may become more solid-like in states of extreme stress, or fluidize to tune protein interactions.

Macromolecular crowding has previously been approximated by a high concentration of molecules of a wide range of sizes (Mourão et al., 2014). Our finding that ribosomal proteins play a dominant role in setting macromolecular crowding adds an interesting caveat to this understanding: the cell *is* crowded by proteins of a large variety of sizes, and there is likely no dominant crowder at the scale of individual proteins, but the crowding environment around large macromolecular complexes is set by ribosome concentration.

It has long been understood that molecular crowding is crucial for biological systems. Our work begins to elucidate why. We show that changes in ribosomal crowding tune phase-separation both *in vitro* and *in vivo.* This finding implies that mTORC1, and indeed any signaling pathway that alters the steady-state concentration of ribosomes, will control the phase-separation of every biological condensate. Thus, our work provides insight relevant to the burgeoning field of phase-separation of cytosolic bio-molecular condensates. Interest in this topic is rapidly growing, as investigators elucidate the impacts of phase-separation on proteins involved in many fundamental processes such as photosynthesis (Freeman Rosenzweig et al., 2017), cell division (Woodruff et al., 2017), development (Brangwynne et al., 2009; Smith et al., 2016), learning (Si et al., 2010), immune signaling (Cai et al., 2014; Hou et al., 2011), and human pathologies including cancer (Kwon et al., 2013), aging and neurodegeneration (Jain and Vale, 2017; Kwon et al., 2014).

**See Supplementary Information for Methods**

## Acknowledgements

We thank Christophe Renou for initial contributions to this project and Monatrice Lam for additional help with FCS measurements. We thank David Savage for initial suggestions about how to build GEMs. We thank David Morgan, David Drubin, Karsten Weis, Jeremy Thorner, Mike Rosen, Amy Gladfelter, Jef Boeke and Douglas Koshland for advice, strains, plasmids and reagents.

We thank William Ludington, Jasna Brujic, Mike Rosen, Jitu Mayor, Ron Vale, Jim Wilhelm, Marcus Taylor, Jan von Skotheim, Josh Zimmerberg for discussions; and Emily Adney, David Truong, Jef Boeke, John Gerhart, Fred Wilt, Ryan Joyner, Carlos Pantoja, Tim Lionnett, Jon Ditlev, Matthew Maurano, Gabrielle Riekhoff, for help with the manuscript.

We acknowledge the help of support of the Janelia Advanced Imaging Center, a facility jointly supported by the Gordon and Betty Moore Foundation and HHMI at HHMI’s Janelia Research Campus in collecting 3D data using the aberration corrected multifocal microscope – data necessary to validate our 2D simplifications.

We gratefully acknowledge funding from the William Bowes Fellows program, the Vilcek Foundation, and the HHMI HCIA summer institute (LJH); the National Science Foundation Graduate Research Fellows Program (GB). Christine Jacobs-Wagner is an Investigator of the Howard Hughes Medical Institute.

## Data Availability

All subtomogram averages presented in this study have been deposited in the Electron Microscopy Data Bank (EMD-XXXX, EMD-XXXX, EMD-XXXX, EMD-XXXX), along with the tomograms from Figure 4 (EMD-XXXX, EMD-XXXX).

## Method Details

### Plasmid construction

The open reading frames encoding the *Pyrococcus furious* encapsulin and *Aquifex aeolicus* (AqLS) lumazine synthase protein based on the published crystal structures (www.rcsb.org 2E0Z and 1NQU respectively) were codon optimized for yeast and mammalian expression and synthesized as IDT gene blocks (www.idtdna.com). The 40nm-GEM plasmid for yeast expression was constructed by fusion at the 5’ with the yeast INO4 promoter and at the 3’ (via a Gly-Ser linker) with the T-Sapphire fluorophore *(Zapata-Hommer and Griesbeck, 2003)* by Gibson assembly into the pRS305 vector (pLH0497: pRS305-PINO4-PfV-GS-Sapphire). The Mammalian expression vector was assembled similarly into the pCDNA3.1 vector (Thermo Fisher) with the CMV2 promoter (pLH611: pCDNA3.1-CMVP2-PfV-GS-Sapphire-GGS). To make a Lentiviral vector (pLH1337: CMV-PfV-Sapphire-IRES-DsRed-WPRE) to express 40nm-GEMs, the PfV-GS-Sapphire sequence was digested from pLH611 and incorporated into a Clontec V4 vector via Gibson assembly. We empirically determined that Sapphire was brighter than GFP in the context of GEMs, presumably because the long Stokes-shift of this fluorophore avoids some of the autoquenching that may occur on the crowded surface of these particles. However, this crowded environment also appears to affect the photochemistry of Sapphire such that fluorophore excitation is efficient at 488 nm thus, for imaging purposes, we used settings optimized for GFP (see Imaging below). The 15nm-GEM for yeast expression was assembled by fusion at the 5’ with the yeast HIS3 promoter and at the 3’ (via a Gly-Ser linker) with the T-Sapphire fluorophore by Gibson assembly into the pRS306 vector (pLH1144:pRS306-PHIS3-AqLumSynth-Sapphire). *μ*NS-GFP (PHIS3-GFP-*μ*NS-URA3) particles were constructed by Gibson assembly of the published N-terminal GFP fusion to the C-terminal fragment of *μ*NS (residues 471-721) *(Broering et al., 2005*) together with the yeast HIS3 promoter into the pRS306 vector (pLH1125: pRS306-PHIS3-GFP-muNS). The Sumo(10)-Sim(6) yeast reporter(pLH1388:pAV106-pTDH3-mCherry-10xSumo-6xSIM) plasmid was generated by chemical synthesis of mCherry fused to a linked Sumo(10)-Sim(6) sequence that was based on the human sequence and then codon optimized for yeast. The mammalian Sumo(10)-Sim(6) was graciously gifted from the lab of Mike Rosen. All yeast plasmids were integrated into the host genome.

### Yeast cell culture and transformation

Yeast strains were created by transforming with a LiAc based approach according to standard methods. BY4741 deletion mutants were obtained from the Yeast Deletion Collection. pLH0497:pRS305-LEU2-PINO4-PfV-GS-Sapphire or pRS306-URA3-PHIS3-PfV-GS-Sapphire was transformed into the collection to allow for screening of mutants or into BY4741 and W303 strains for the rest of the experiments. The cdc28-as1 strain was taken from *(Bishop et al., 2000)*. A list of yeast strains constructed is provided in Table S2 and STAR methods. Exponentially growing cultures O.D. ≥ 0.1 and ≤ 0.4 were used in all experiments unless otherwise noted. Note: It is extremely important to avoid culture saturation - all cultures were started from single colonies and grown overnight to log phase (typically we set up 1/5 serial dilutions to catch one culture at the correct OD). If cultures saturate, it takes many generations to reset the cellular rheology. All strains were grown at 30^o^C in a rotating incubator.

### Mammalian cell culture conditions and virus production

Mammalian cells were maintained at 37^o^C with 5% CO_2_. HEK293 and HEK293T were grown in high glucose DMEM (Life Technologies) supplemented with 10% fetal bovine serum (FBS; Gemini Bio-products), penicillin (50 U ml^-1^) and streptomycin (0.05mg.ml^-1^) (Life Technologies) unless otherwise stated. In order to create lentivirus, 800,000 HEK293T cells were plated in 10mL media in 15cm dishes. The next day, each well was transfected with 24*μ*g vector, 1.2*μ*g tat, 1,2 **μ*g* rev, 1,2*μ*g gag/pol, and 2.4*μ*g of vsv-g DNA with 90*μ*L trans-IT in 2mL DMEM. Supernatants were collected at 24, 48, and 72 hours after transfection and stored at 4^o^C until they were spun at 16.5K for 90 minutes on a Beckman L-80 Ultracentrifuge. Viral pellets were resuspended in 1/50th of their original volume in DMEM (with 10% FBS) and stored at -80^o^C until their use. Stable HEK293 cell lines were created by transfection with (pLH611: pCDNA3.1-CMVP2-PfV-GS-Sapphire-GGS) followed by neomycin selection. Additional HEK293 cell lines were created by lentiviral transduction with pLH1337-CMV-PfV-Sapphire-IRES-DsRed-WPRE. No differences in terms of cellular rheology were seen between these different methods. In order to transduce these cell lines, 50,000 cells were plated in 2mL of media in 6 well plates. The next day, media was removed and replaced with media containing 8 *μ*g/mL polybrene. Between 1-20*μ*L of concentrated virus was added to the well and then the media was replaced after 24 hours.

### Drug treatment

In order to inhibit mTORC1 signaling, we treated with rapamycin (Tocris Bioscience, Avonmouth, Bristol, UK) at 1*μ*M from a 1mM stock. In order to block ribosome production, we treated with PolI inhibitors BMH21 and CX5461 (Selleckchem, Houston, Texas, USA) at concentrations of 10uM and 500nM, from stocks of 50mM and 1mM, respectively. In order to increase autophagy, we treated with SMER28 (Tocris Bioscience, Avonmouth, Bristol, UK) at a concentration of 5uM from a stock of 10mM. In order to decrease autophagy, we treated with Wortmannin (Cell Signaling Technology, Danvers, MA, USA) at a concentration of 800nM from a stock of 4mM. All stocks were prepared in DMSO and stored at -20^o^C until needed. DMSO was used as an vehicle control in all experiments. In order to de-polymerize actin and microtubules in yeast we used 200*μ*M latrunculinA and 15*μ*g/mL nocodazole (Tocris). In order to depolymerize actin in HEK293 cells we used 4*μ*g/mL latrunculin (tocris) A and in order to freeze the cytoskeleton we used 1mM jasplakinolide (Cayman), 1mM latrunculinA (Tocris) and 5mM y27632 (selleckchem) *(Peng et al., 2011).*

### FCS and coefficient of diffusion of 2xGFP

A custom-modified inverted microscope (Nikon Eclipse Ti; Nikon Instruments) was used for FCS measurements. Prior to each measurement, a focus spot within a cell was located by eGFP epiflu-orescence. A 100-ps pulsed 482 nm diode laser (PicoQuant) was coupled to a single-mode fiber and collimated to a 4-mm diameter, then focused on the sample through a 100x objective (CFI Apo 100x Oil immersion TIRF NA 1.49; Nikon Instruments), with the laser power of 0.2 *μ*W before the objective. The focus spot was calibrated with a fluorescent dye with a known diffusion coefficient (Alexa 488, D = 435 *μ*m^2^/s *(Petrášek and Schwille, 2008)).* Each FCS measurement was the average of 10-20 cells. Fluorescence emitted from the sample was passed through a 50-*μ*m pinhole (Thorlabs), and focused to a bandpass-filtered single-photon avalanche diode with a 150 x 150 *μ*m element (PDM module; Optoelectronic Components). The resulting fluorescence fluctuation was processed by a hardware correlator (Correlator.com), which generated the autocorrelation function. See Figure S1 for results and more details on the fitting procedure of the autocorrelation function.

### Imaging and direct particle tracking

Single particle tracking in *Saccharomyces cerevisiae* was performed for the 40nm-GEMs, AqLS particles, RNA particles, and *μ*NS. The particles were imaged using TIRF Nikon TI Eclipse microscope at 488nm, and their fluorescence was recorded with a scMOS (Zyla, Andor) with a 100x objective (pixel size 0.092*μ*m), with a time step that depends on the particles. The GEMs were imaged at a rate of one image every 10ms, whereas both the RNA particles and the *μ*NS were imaged at 100ms time step.

Single particle tracking in HEK293 cells was performed for 40nm GEMs using an Andor Yokogawa CSU-X confocal spinning disc on a Nikon TI Eclipse microscope and their fluorescence was recorded with a sCMOS Prime95B camera (Photometrics) with a 100x objective (pixel size 0.1*μ*m), at 10ms image capture rate.

The tracking of particles was realized through the Mosaic suite of FIJI, using the following typical parameters: radius = 3, cutoff = 3, 10% of intensity of fluorescence, a link range of 1, and a maximum displacement of 8 px, assuming Brownian dynamics.

### Extraction of the rheological parameters

Various parameters were extracted from the trajectories of particles. For every trajectory, we calculated the time-averaged mean-square displacement (MSD), as defined in *(Munder et al., 2016)*, as well as the ensemble-average of the time-averaged MSD. As observed in the insets figure 2D and 2E, where the ensemble-averaged MSD is plotted as a function of time in a log-log plot, the diffusion of the tracer particle is subdiffusive, and generally obeys the following law:

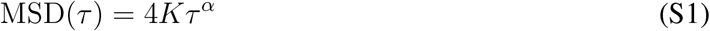

where *α* is the power exponent of the anomalous diffusion, *α* ≤ 1 in the case of a subdiffusive behavior. In this case, the apparent diffusion coefficient, *K*, is not in units of **μ**m^2^/s, but rather in units of **μ**m^2^/s^*α*^.

To characterize individual particle trajectories, we simplified to a linear MSD fit to measure an effective diffusion at short time scales (less than 100 ms for GEMs, 1s for mRNP and **μ**NS particles). To do this, we calculated the MSD and truncated it to the first 10 points, and fitted it with the following linear relationship:

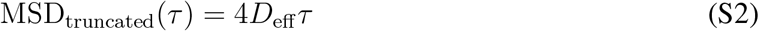

where *D*_eff_ is the effective coefficient of diffusion of the tracer particle, and plotted the distribution of this effective diffusion coefficient. We use the Kolmogorov-Smirnov statistical test to assess the statistical difference between distributions (kstest2 function in Matlab).

### EM grid preparation and data acquisition

W303 *S. cerevisiae* were grown on YPD plates for two days, then suspended in SCD media at low cell concentration by serial dilution and grown overnight at 30^o^ C on a roller drum to an OD of 0.25. Cells were then incubated with 1*μ*M rapamycin in DMSO or only DMSO (control) for 2 h until an OD of 0.55. Cells were frozen onto EM grids from 2-2.5 h after addition of the drug. 4 *μ*L of culture was applied to R2/1 holey carbon copper EM grids (Quantifoil) and immediately vitrified by plunge-freezing into a liquid ethane/propane mixture with a Vitrobot Mark IV (FEI, The Netherlands) using a blot time of 10 s, a blot force of 10, and a chamber conditioned to 25^o^C and 90% humidity. EM grids with vitrified yeast cells were transferred either to a Quanta or Scios dual-beam microscope (both FEI, The Netherlands) for focused ion beam micromachining. The vitrified cells were platinum coated with organometallic platinum and subsequently thinned by scanning gallium ions in a stepwise fashion from both sides. This yielded vitrified cellular sections of 100-200 nm thickness that were suitable for cryo-electron tomography *(Schaffer et al., 2015, Schaffer et al., 2017*). EM grids with milled samples were transferred to a Titan Krios TEM (FEI, The Netherlands) operated at an acceleration voltage of 300 kV, an object pixel size of 3.42 *A* and a nominal defocus of -6 *μ*m. The TEM was equipped with a Quantum energy filter (Gatan) and a K2 summit direct electron detector (Gatan) operated in movie mode (12 frames per second). Single-axis tilt series were acquired in SerialEM *(Mastronarde, 2005)* using a bi-directional tilt scheme covering a tilting range of approximately -60^o^ to 60^o^ with a 2^o^ angular increment. Depending on the pre-tilt of cellular sections in the TEM, the two tomogram halves were connected at either +20^o^ or -20^o^ tilt. The cumulative electron dose for a tilt series was 70-120 electrons per A^2^, depending on the sample thickness.

### Tomogram reconstruction

Frames from the K2 direct detector were aligned with MotionCor2 (*Zheng* et al., *2017*) using 3x3 patches for local alignment. For each tilt series, the resulting frame-aligned projections were sorted according to their tilt angles and compiled into an image stack that was loaded into IMOD for tilt series alignment via patch tracking. Projection-wise translations and rotations determined during patch tracking were extracted from IMOD’s output files and used for tilt series alignment in TOM/AV3 (*Nickell* et al., *2005*, *Förster and Hegerl, 2007*). Phase reversals introduced by the contrast transfer function (CTF) were determined on each individual projection using strip-based periodogram averaging *(Eibauer et al., 2012)* in TOM/AV3 and corrected in PyTom *(Hrabe et al., 2012).* Finally, the aligned CTF-corrected tilt series was weighted for subsequent reconstruction of tomographic volumes via weighted back projection (AV3/TOM). For reconstruction of binned tomograms, the tilt series was scaled to 2.1 nm in Fourier space (AV3/TOM).

### Determination of the cytosolic volume

Binary masks encompassing exclusively the cytosolic volume were generated by manual segmentation of tomograms in Amira (FEI, The Netherlands). As each voxel corresponds to a volume of (2.1 nm)^3^ = 9.26 nm^3^, the exact cytosolic volume included within the tomogram could be obtained by counting the voxels encompassed by the mask.

### Subtomogram analysis

A) Ribosome: To generate a data-driven *de novo* template for correlation-based ribosome localization, 500 ribosomes were manually selected from one of the tomograms and reconstructed as described below. The subtomograms were iteratively aligned using fast rotational matching (FRM) *(Chen et al., 2013*) implemented in PyTom with a featureless sphere as a starting reference (Figure S2A). The average converged into a ribosome within 12 iterations and was subsequently used as a template for correlation based localization of ribosomes (*Frangakis* et al., *2002*) in all tomograms. For each tomogram, the cross-correlation function resulting from template matching was masked to include only the cytosolic volume of the cell (Figure S2B) and the 5000 highest-scoring peaks were extracted. To avoid multiple detection events for the same ribosome, a minimal Euclidean distance of 18.9 nm (9 voxels) between peaks was imposed. The distribution of correlation coefficients for the extracted peaks showed clear separation of coefficients corresponding to true and false positives (Figure S3). This allowed fitting of a Gaussian function to the distribution of coefficients corresponding to true positives and thus quantification of ribosome abundance within the cytosolic volume.

For detailed analysis of ribosome structures, all ribosomal particles with correlation coefficients better than one standard deviation below the mean of the fitted Gaussian function were retained and reconstructed at full spatial resolution in PyTom from the CTF-corrected, weighted and aligned projections covering approximately the first half of the tilt series. Projections corresponding to the second half of the tilt series were excluded at this step due to excessive beam damage that dampens high-resolution signal. The reconstructed subtomograms were aligned until convergence with Relions gold standard 3D auto-refine functionality, which is now available for subtomograms *(Bharat et al., 2015*). During subtomogram averaging, Relions 3D CTF model was used to compensate for beam damage with the recommended B-factor of -4 per electron per Å^2^. Resolution of the resulting averages was estimated based on Fourier shell cross-correlation (FSC) of two completely independent halves of the data using FSC = 0.143 as the cutoff criterion. For computation of the difference density between ribosome structures from control and rapamycin-treated cells, the averages were filtered to 15 Åresolution, normalized according to density mean and density standard deviation, and subtracted from each other. The UCSF Chimera software package *(Goddard et al., 2007)* was used for visualization of EM densities.

B) GEMs: GEMs are readily visible in tomograms as high-contrast sphere-like particles (Figure S3D). Consequently, template matching against a hollow sphere of appropriate size in combination with visual inspection of the 50 highest scoring cross-correlation peaks in the cytosolic volume allowed highly specific localization of GEMs in the tomograms. Subtomogram reconstruction, alignment and resolution estimation were performed as described for the ribosome, with the only exception that icosahedral symmetry was applied during subtomogram alignment.

### Osmotic perturbation experiments and cell volume measurement

In order to calculate the dependence of the volume fraction of crowding agent on diffusion of GEMs, we performed hyper- and hypo-osmotic stresses (see model below). LH2129 (BY4741 + PINO4::PINO4-PfV-GS-Sapphire-LEU2) cells were grown in log phase to an OD of 0.3, then spun down for 1 minute at 10000 rpm. Cells were washed with fresh medium, and placed in synthetic complete with dextrose medium complemented with 0M, 0.5M, 1M, 1.5M or 2M of sorbitol. A subset of cells were directly (within 15 minutes) imaged for diffusion, and phase pictures were taken in order to assess cell area as a proxy for cell volume. The rest of the cells were left at various ODs in a shaker at 30^o^C to adapt to the osmotic stress and grow overnight. The next day, cells were imaged for diffusion and cell volume (green points (Figure S8), to check that cell volume and the diffusion of particles had recovered to their nominal values. These pre-adapted cells (which have built up a high concentration of internal osmolyte) were then spun down and placed in regular CSM, creating a hypo-osmotic stress of -0.5M, -1M, -1.5M and -2M, and immediately imaged for diffusion and cell volume. The same process was used for HEK293 cells, with osmotic stress of 0.25M and 0.5M sorbitol. Cells were trypsinized and their volume measured from their area when the cells were spherical.

### TSC1 siRNA experiments

TSC1 (s14433 or s14434) was targeted by Silencer Select siRNAs from Thermo Fisher Scientific. 75 pmoles of siRNA were transfected using Lipofectamine RNAiMAX transfection reagent from Thermo Fisher Scientific as per manufacturer’s instructions. Cells containing GEMs were assayed for diffusion at 72 hours post transfection. Knockdown was validated by western blot using Hamartin (TSC1) (D43E2) Rabbit mAb #6935 from Cell Signaling Technologies using standard techniques.

### Protein purification

Proteins were expressed in Rosetta2 DE3 competent cells by induction with 100 M IPTG for 18 hr at 16C. 4 liters of bacterial culture were collected and centrifuged at 4000rpm for 20 min at 4C. The cell pellet was resuspended in 100 ml cold lysis buffer (50mM NaH2PO4, 300mM NaCl, 10 mM imidazole pH7.6) containing 1 mM PMSF. After sonication, the lysate was centrifuged at 12000rpm for 30 min at 4C. The supernatant was mixed with 8 ml of 50% slurry of Ni-NTA beads (Qiagen). The lysate was incubated with beads for 2 hour at 4C. The bound beads were collected by centrifugation at 500g for 1 minute and rinsed 3 with 30 ml bacterial wash buffer containing (50mM NaH2PO4, 300mM NaCl, 20 mM imidazole pH7.6). The bound proteins were eluted with 8ml elution buffer (50mM NaH2PO4, 300mM NaCl, 500 mM imidazole pH7.6).

The elution was exchanged into 2 ml of SUMO-SIM protein buffer (150mM KCl/20mM HEPES pH7/1mM MgCl2/1mM EGTA/1mM DTT) using a PD10 column (GE Healthcare), followed by further concentrating to 300-600 M with Amicon Ultra 30K device (Millipore) at 4C.

SIM is tagged with Alexa Fluor 488, not fused with GFP. The protein was conjugated with Alexa Fluor 488 with large scale antibody/protein labeling Kits (A10235, Thermo Scientific).

### Phase separation experiments

In order to determine if ribosomes are capable of acting as a crowding agent *in vitro*, we added purified ribosomes from an *in vitro* translation kit (IVT) (NEB, Ipswich, MA) to a mix of purified Small Ubiquitin like Modifier (SUMO) 60 *μ*M module and SUMO Interaction Motif-GFP (SIM-Alexa Fluor 488) 60*μ*M module. Ribosomes were added at the same concentrations measured *in vivo* by cryo-ET as well as at an intermediate concentration. Ribosomes, SUMO, and SIM were mixed in a well of a 384 well imaging plate, the top was then covered with clear tape and then the plate was allowed to sit overnight in order to reach a steady-state before imaging. The plate was imaged on an Andor Yokogawa CSU-X confocal spinning disc on a Nikon TI Eclipse microscope and GFP fluorescence was recorded with an scMOS (Zyla, Andor) camera with a 100x objective (pixel size 0.1*μ*m). Images were loaded in FIJI and the partition coefficient (amount of protein that has condensed into liquid droplets versus protein dissolved in the bulk aqueous phase) was calculated by segmenting the image into two categories: bright droplets and background. Then the total amount of fluorescent protein was measured in each category through using the raw integrated density value. The partition coefficient was taken as the ratio of protein in the condensed phase versus the bulk phase and plotted in MATLAB.

In order to determine the effects of changes of ribosome concentration via mTORC1 signaling on phase separation, we expressed a mCherry-SUMO(10)-SIM(6) fusion protein in yeast (pLH1388) and mammalian cells (pLH 1389). WT and mutant yeast cells were grown overnight to log phase and then treated with rapamycin for 2 hours. Sorbitol was added in the last ten minutes in indicated conditions. Mammalian cells were treated for 3 hours with rapamycin with sorbitol added in the final 30 minutes where indicated. TIRF microscopy on a Nikon-TI microscope was performed using a 561 nm laser sample through a 100x objective (CFI Apo 100x Oil immersion TIRF NA 1.49; Nikon Instruments). Images were segmented in FIJI to determine the 1) average size of droplets, 2) number of droplets and 3) number of cells. Using these data we then found the total phase separated area as:

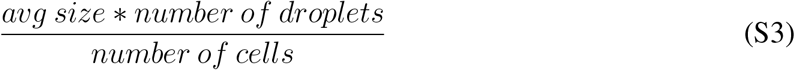

### Fluorescence Correlation Spectroscopy Calculations

FCS data were fitted using a “blinking and anomalous diffusion” model, that has the following form (*Brazda et al., 2011*):

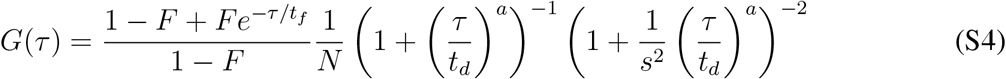

The term on the left before the 1/*N* is the blinking term corresponding to the properties of GFP. In this term, *t_f_* was measured independently from whole cell lysate, *t_f_* = 3.5xl0^-5^s. The term on the right corresponds to the anomalous 3D diffusion of GFP, where *t_d_* is the particle residence time in the focus volume, *t_d_* = *w*^2^/4D. *w* = 220nm and s/w are the radial and axial dimensions of the 3D Gaussian laser focus, respectively, and they were measured using a dye with a known diffusion coefficient (Alexa Fluor 488). In practical terms, *s* does not affect the fit, and was fixed to be *s* = 10. The result of the fit is summarized in table S1, and yields D_DMSO_ = 13.3 ± 1.3 **μ*m*^2^/s and D_rapamycin_ = 12.2 ± 2.8 **μ*m*^2^/s, which are not significantly different (3 biological replicates, n ≥ 10 cells per condition). Note that the anomalous diffusion exponent *a* is in both cases ~ 0.8, close to the measured anomalous parameter measured for GEMs. We chose an anomalous model, as commonly used in the literature to describe motion of GFP in a cell (*Slaughter et al., 2007*), and as it yielded a better fit than the normal diffusion model.

**Table S1:** Results from fitting the blinking and anomalous diffusion model.

| Parameter | DMSO condition | Rapamycin condition |
| --- | --- | --- |
| $N$ | $1.22 \pm 0.03$ | $1.49 \pm 0.09$ |
| $F$ | $0.16 \pm 0$ | $0.29 \pm 0.05$ |
| $t_f$ (s) | $3.5 \times 10^{-5}$ (fixed) | $3.5 \times 10^{-5}$ (fixed) |
| $t_d$ (s) | $(9.1 \pm 0.9) \cdot 10^{-4}$ | $(9.9 \pm 2.3) \cdot 10^{-4}$ |
| $s$ | 10 (fixed) | 10 (fixed) |
| $a$ | $0.79 \pm 0.05$ | $0.77 \pm 0.07$ |

### Crowding regulation through control of ribosome concentration

#### Model Basis

In the following, we derive a model of crowding control in the cell. The purpose of the model is to link cell volume change to changes in the diffusion of a tracer particle, like our 40nm-GEMs. We assume that there is a major source of crowding within the cell, that is impacting the diffusion coefficient of 40nm-GEMs. We express the diffusion of the tracer particle as a function of volume fraction of the major crowder source, φ, using the phenomenological Doolittle equation *(Doolittle, 1951, Hunter and Weeks, 2012*):

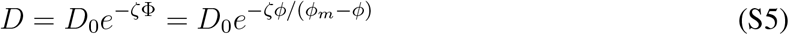

where *D*_0_ is the coefficient of diffusion in an infinitely diluted solution, *φ_m_* the maximum fraction of the crowder, and *ζ* is a constant. We write the volume fraction for the major source of crowder as:

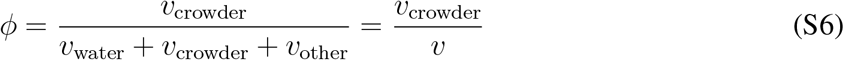

where *υ*_crowder_ is the volume occupied by the major source of crowding, *υ*_other_ the volume occupied by other macromolecules, and *υ*_water_ the volume occupied by water in the cell. *υ* is the volume of the cell. The maximum fraction of crowder in the cell is reached when the volume of water is close to 0, such that *φ_m_* ~ *υ*_crowder_/(*υ*_crowder_ + *υ*_other_).

#### Validation of Doolittle equation and determination of parameters using instantaneous volume change through osmotic stress

During an instantaneous volume change as a result of an osmotic stress, the total number of macromolecules remains, to a first approximation, constant. The cell volume changes because of a passive outflow (hyper-osmotic stress) or inflow (hypo-osmotic stress) of water. We denote φ_0_ the volume fraction of macromolecules before the osmotic shock, when the cell volume is *υ*_0_:

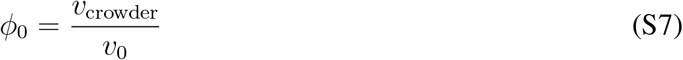

Denoting 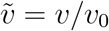 the normalized cell volume, one can express Φ:

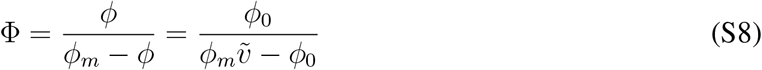

Note that the diffusion coefficient *D*_0_ in equation S5 does not correspond to the coefficient of diffusion under normal conditions, but corresponds to the coefficient of diffusion for an infinitely diluted solution of macromolecules. Rather, the coefficient of diffusion under normal conditions, that we denote 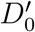, is defined when Φ = Φ_0_ = *φ*_0_/(*φ_m_ – φ*_0_):

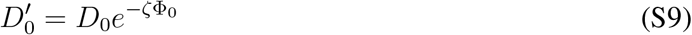

which leads to the formula that describes the instantaneous change of the coefficient of diffusion upon a given volume change 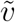:

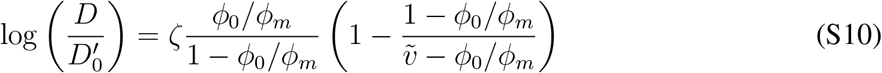

We used equation S10 to fit the coefficient of diffusion of 40nm-GEMs under hypo- and hyperosmotic stresses (see figure S8). The model is in good agreement with our data (r^2^ = 0.85), and gives parameters for *S. cerevisiae ζ* = 0.61 ± 0.2, *φ*_0_/*φ_m_* = 0.54±0.5.

This number means that, under normal conditions, that the fraction of crowder inside the cell is about 50% the maximum crowding. This number is the non-osmotic volume, *φ*_0_/*φ_m_* = *υ*\*, which corresponds to the volume of the cell occupied by macromolecules *(Miermont et al., 2013*).

We performed the same osmotic stress experiment on HEK293 cells, and initially measured different parameters (ζ ~ 1.6, *υ*\* ~ 0.35). Osmotic stress is known to strongly affect the actin cytoskeleton in mammalian cells, which could affect the interaction parameter, *ζ*, which was confirmed when we treated the cells with LatA at the same time we did the osmotic stress (ζ ~ 3.6, *υ*\* ~ 0.35): the interaction parameter of the GEMs with the environment increased. When the actin cytoskeleton was stabilized with JLY cocktail, we found that the 2 parameters of the model were closer to the yeast values: ζ ~ 0.6, is very similar to yeast, suggesting that the interactions of the GEMs with the microenvironment is the same, and *υ*\* ~ 0.35 is lower, suggesting that mammalian cells are less crowded.

This result suggest that 40nm-GEMs seem may interact with the same species inside both *S. cerevisiae* and HEK293 cells in a diffusion dependent manner.

#### Homeostatic crowding, and homeostasis breaking under a rapamycin treatment

What is the major source of crowding in the cell?

Our mutagenesis experiments suggested that mTORC1 controls cytosolic fluidity by tuning ribosome biogenesis and degradation. Therefore, ribosome concentration, and the concentration of proteins obtained through translation are candidates for the major source of crowding. We blocked translation using cycloheximide and found that the diffusion coefficient of 40nm-GEMs was not affected treatment (figure S1). This result suggests that ribosomes are the most important source of crowding regulation for the 40nm-GEMs.

Therefore, we can re-write the volume fraction of crowder, considering ribosomes as the major crowder:

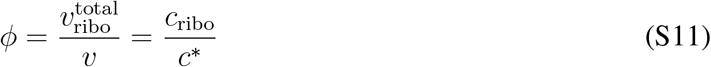

with *c*\* *=* 1/*υ*_ribo_, *υ*_ribo_ being the typical volume of a single ribosome.

This leads to the following equation:

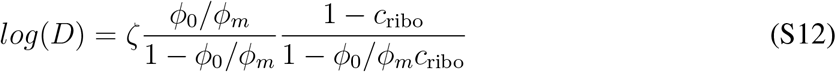

We used this equation with the parameters measured by an osmotic stress to predict how particular mutations or chemical treatment should affect crowding, measured through the coefficient of diffusion, as a function of ribosome concentration. The ribosome concentration is determined by its number in the cells, N, and the volume of the cell, v. We measured the number of ribosomes N either by direct counting in EM, or their relative amount to wild-type or normal conditions was assessed by quantification of a total nucleic acid extraction run on an agarose gel (see Fig S8E-G). The cell volume was determined through brightfield measurements.

Fig. 4 displays the model prediction for both *S. cerevisiae* and HEK293 cells, which is in very good agreement with the measured data. This suggests that:

- Ribosomes are indeed the main determinant of cytosolic crowding inside the cell and can be considered as hard spheres.
- The cytoplasm of mammalian cells and yeasts behave similarly in terms of crowding.

## Supplementary movies

### Supplementary movie 1

Bright field images (top panels), fluorescent images (middle panels) and fluorescent images with single particle trajectories superimposed (bottom panels), for DMSO (vehicle control, left) and rapamycin treated (right) *S. cerevisiae.* Scale bars are 5*μ*m. The movie playback speed is set to 100 frames per second (roughly real time), and the average time between frames is 8 ms. The time stamp shows seconds.

### Supplementary movie 2

Fluorescent images (top panels) and fluorescent images with single particle trajectories superimposed (bottom panels), for DMSO (vehicle control, left) and rapamycin treated (right) HEK293 cells. Scale bars are 4μm. The movie playback speed is set to 100 frames per second (roughly real time), and the average time between frames is 8 ms. The time stamp shows seconds.

### Supplementary movie 3

In situ cryo-ET of DMSO-treated control yeast cells. Related to figure 4A. The movie slices back and forth in Z through the tomographic volume, then reveals the 3D segmentation of the non-cytosolic volume (grey) and structures of the 40nm-GEMs (magenta), and then reveals the structures of the ribosomes (cyan).

### Supplementary movie 4

In situ cryo-ET of rapamycin-treated yeast cells. Related to figure 4B. The movie slices back and forth in Z through the tomographic volume, then reveals the 3D segmentation of the non-cytosolic volume (grey; aggregates in dark grey) and structures of the 40nm-GEMs (magenta), and then reveals the structures of the ribosomes (cyan).

### Supplemental Figures

**Figure S1:**
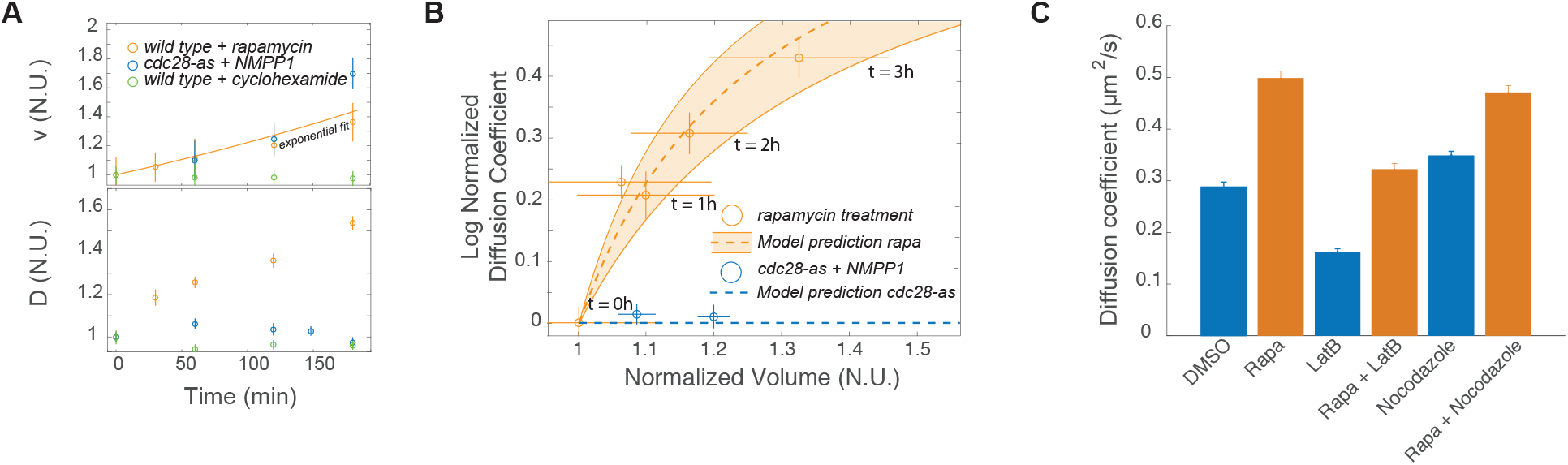
Cell volume increase, translation, and cytoskeletal pertubations do not explain rapamycin-dependent effects on GEM behavior. Related to figures 2 and 3. (**A**) Change in normalized volume and coefficient of diffusion over time for rapamycin treatment (orange), inhibition of cell cycle in conditional mutant cdc28-as allele background with NMPP1 (blue) and cyclohexamide treatment (green). (**B**) log normalized diffusion coefficient can be predicted by our model of ribosomal crowding and cell volume increase. (**C**) Latrunculin and Nocodazole treatment alter the diffusion coefficient of GEMs but do not prevent the increase in diffusion rate under rapamycin treatment.

**Figure S2:**
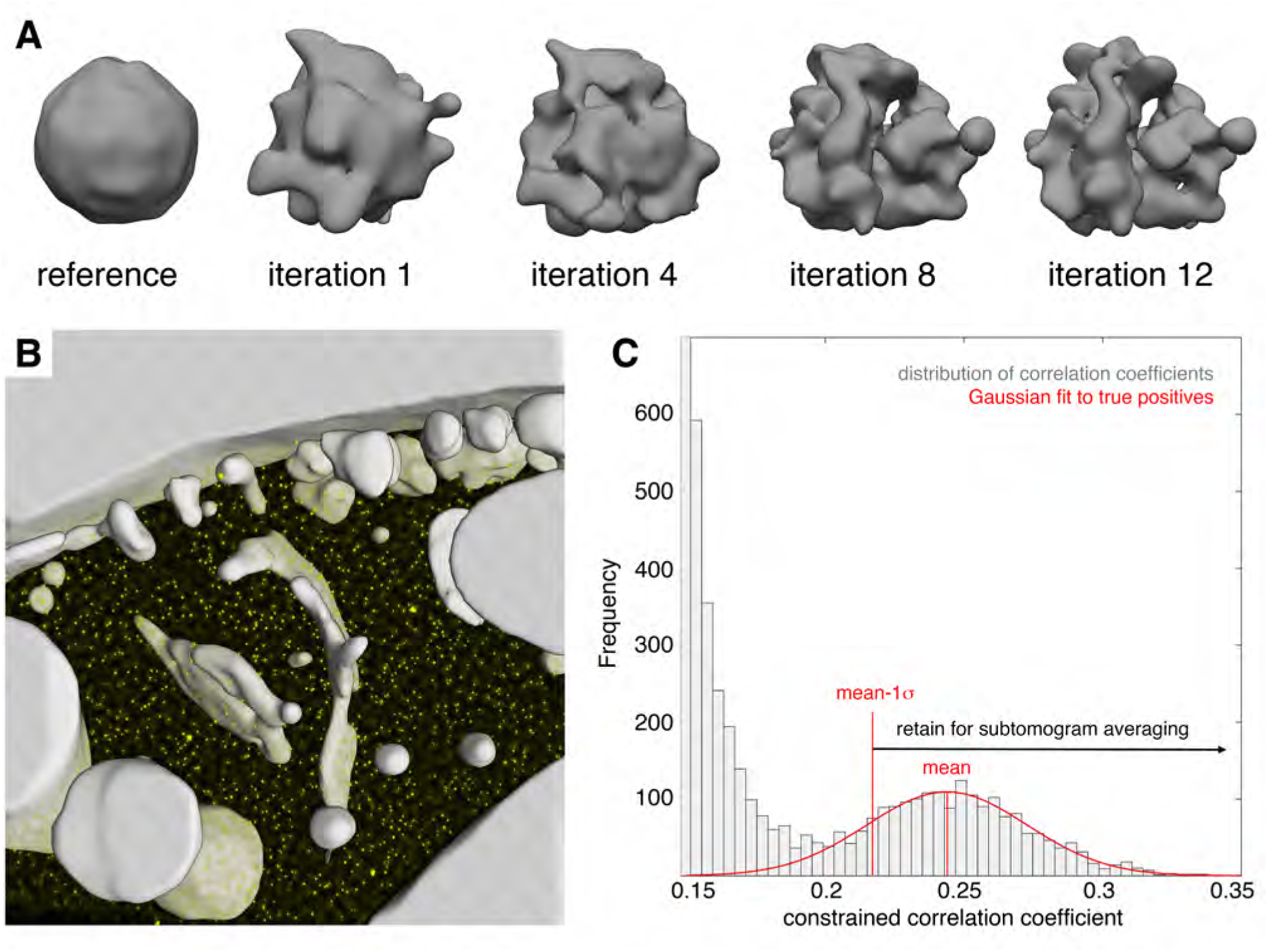
Ribosome quantification via cryo-electron tomography and template matching. Related to figures 4 and 6. (**A**) 500 manually selected ribosome-containing subtomograms were iteratively aligned with a sphere as a starting structure (left). Within 12 iterations, the averaged density converged to a yeast 80S ribosome (right) that was subsequently used as a purely data-driven *de novo* template for correlation-based ribosome localization (template matching) in the tomograms. (**B**) Example cross correlation function (yellow) obtained from template matching against the *de novo* ribosome structure, superposed with the non-cytosolic cellular volume (gray) excluded from the analysis. Peaks in the cross-correlation function (yellow spots) indicate putative ribosome positions. (**C**) Distribution of cross-correlation coefficients for the 5000 highest-scoring peaks, which were extracted from the cross-correlation volume depicted in B while imposing a minimal Euclidean distance of 18.9nm (9 voxels) between peaks. A Gaussian function (red) was fit to the distribution of coefficients corresponding to true positives. The integral of the Gaussian function corresponds to the number of ribosomes included in the cytosolic volume.

**Figure S3:**
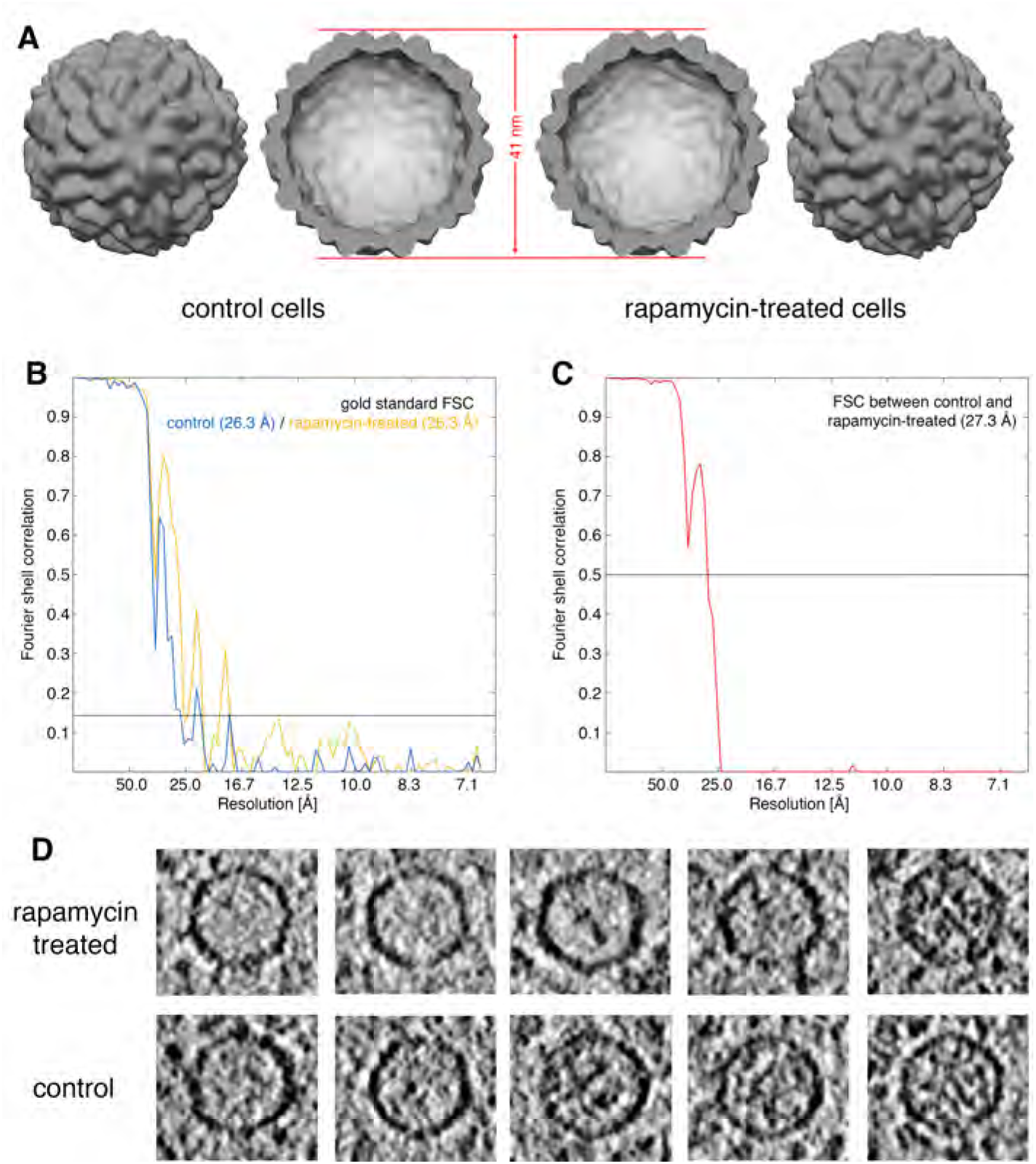
GEM structures from control and rapamycin-treated cells are indistinguishable. Related to figures 1 and 4. (**A**) 30nm-GEM subtomogram averages obtained for control (left panels) and rapamycin-treated (right panels) cells filtered to 26.3 Å resolution. In the central two panels, averages have been sliced open to show the interiors. (**B**) Fourier shell correlation (FSC) between subtomogram averages derived from two independent halves of the dataset (gold standard) for control (blue) and rapamycin-treated (orange) cells. Resolution was determined to be 26.3 A in both cases using the FSC = 0.143 resolution criterion. (**C**) FSC between the two resolution-limited subtomogram averages obtained for control and rapamycin-treated cells. High correlation (FSC > 0.5) within the trustworthy resolution range suggests that GEM structures under both conditions are identical. (**D**) Gallery of individual GEM particles from control (lower row) and rapamycin-treated (upper row) cells. Each image corresponds to a central tomogram slice through the GEM particle. The amount of cargo within the GEM lumen varies.

**Figure S4:**
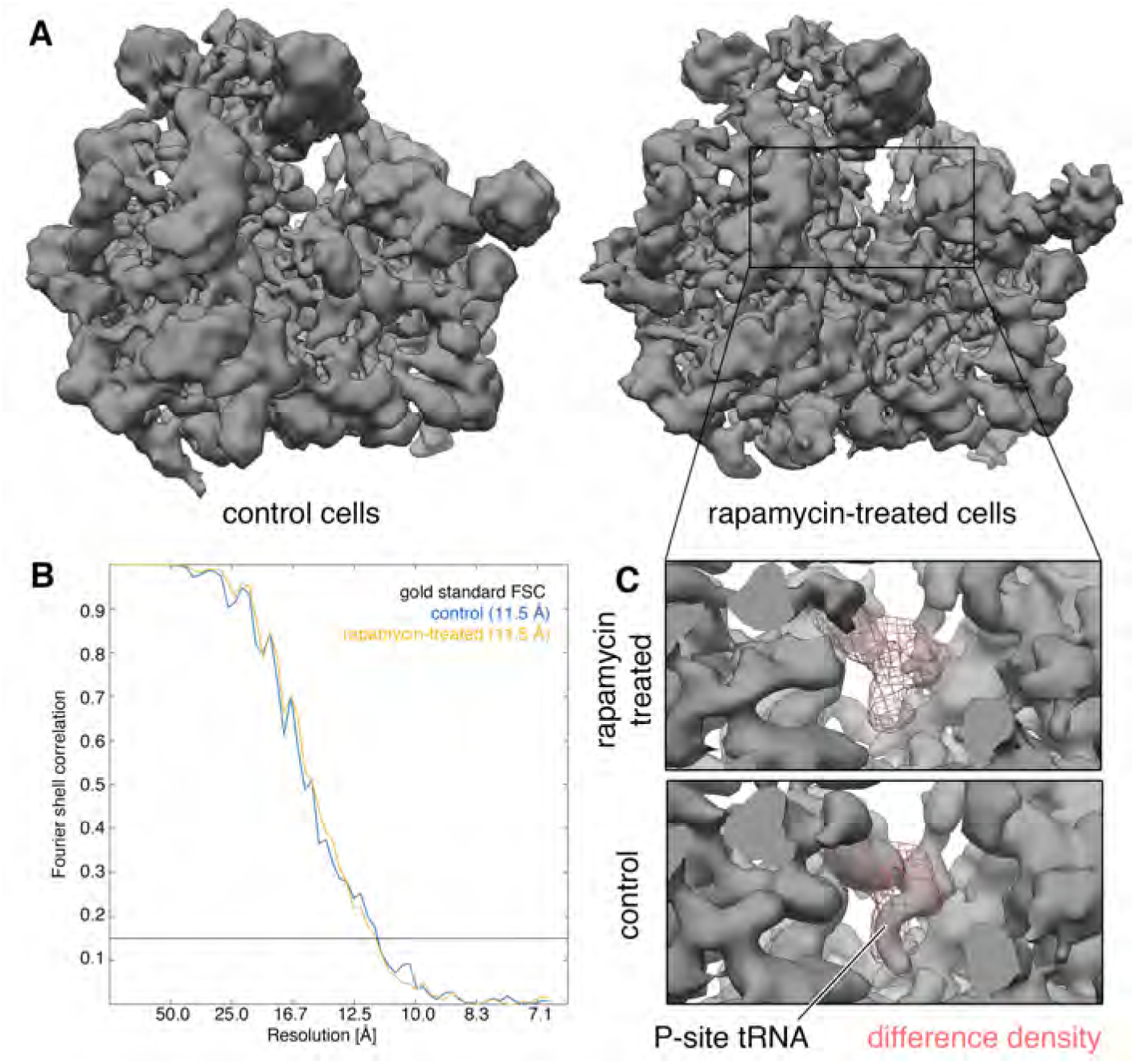
Ribosome structures from control and rapamycin-treated cells differ in P-site tRNA density. Related to figure 4. (**A**) Ribosome subtomogram averages obtained for control (left) and rapamycin-treated (right) cells filtered to 11.5 Åresolution. (**B**) FSC between subtomogram averages derived from two independent halves of the data (gold standard) for control (blue) and rapamycin-treated (orange) cells. Resolution was determined to 11.5 Åin both cases using the FSC = 0.143 resolution criterion. (**C**) Enlarged view of the region indicated with a box in **A**, comparing the ribosome structures from rapamycin-treated (upper panel) and control (lower panel) cells. The most significant density difference between both ribosome structures co-localizes with the P-site tRNA, which is resolved in the control but not the rapamycin-treated condition.

**Figure S5:**
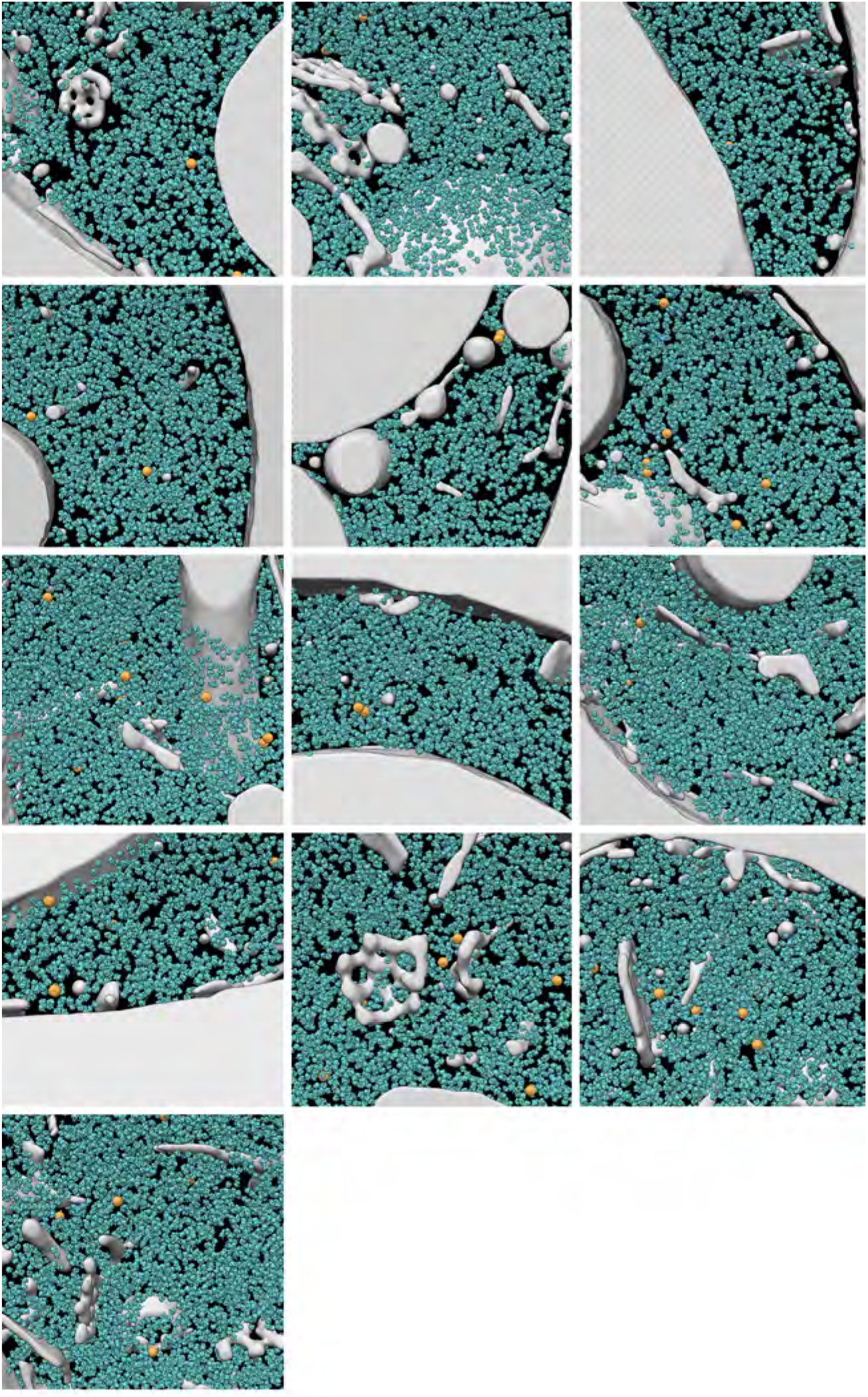
Gallery of 3D segmentations from the complete cryo-ET dataset of DMSO-treated control yeast cells. Related to figure 4A. Detected ribosomes are depicted in blue, GEMs in orange and the non-cytosolic volume that was excluded from the analysis in gray. The example tomogram in Fig. 3A is not pictured here.

**Figure S6:**
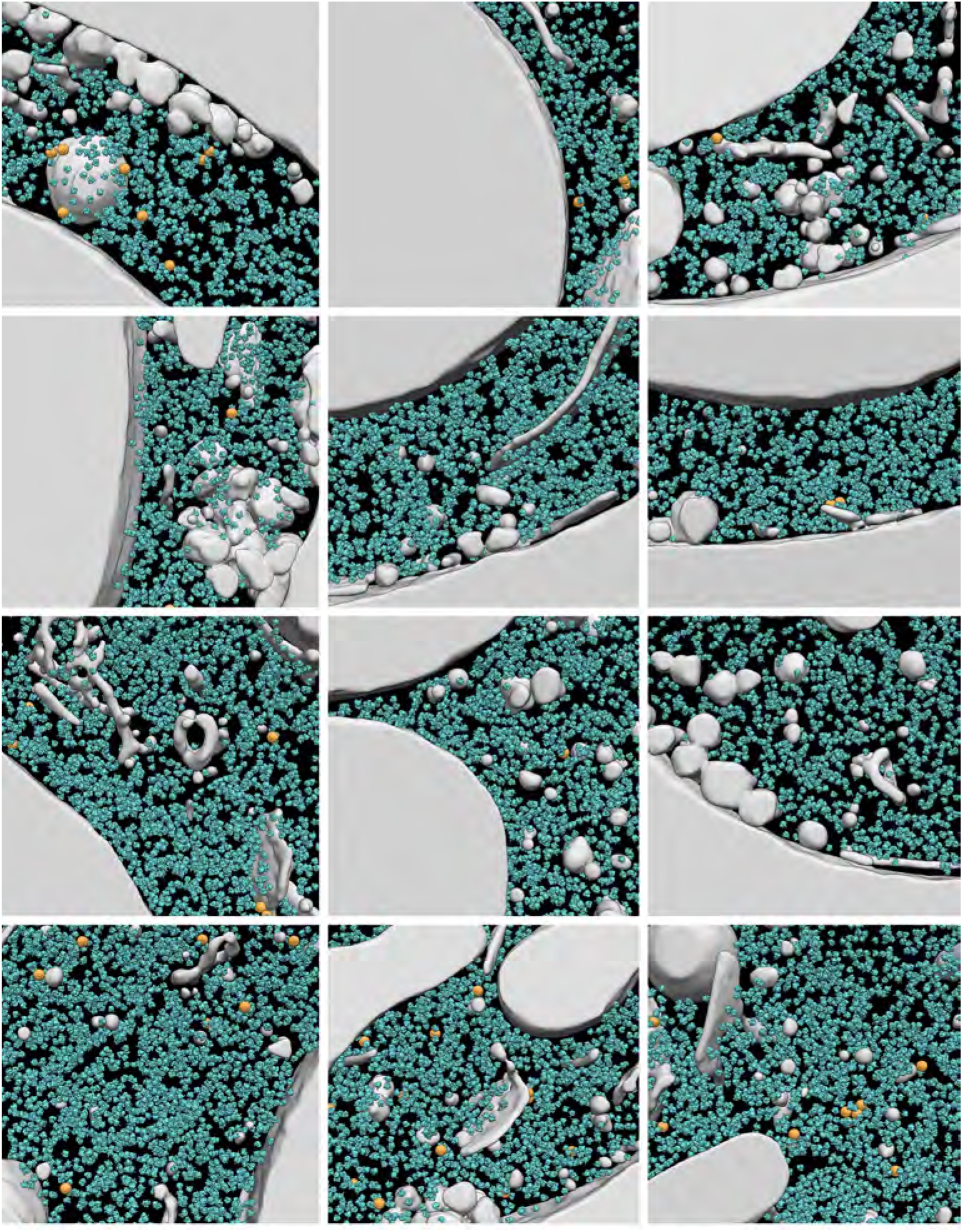
Gallery of 3D segmentations from the complete cryo-ET dataset of rapamycin-treated yeast cells. Related to figure 4B. Detected ribosomes are depicted in blue, GEMs in orange and the non-cytosolic volume that was excluded from the analysis in gray. The example tomogram in Fig. 3B is not pictured here.

**Figure S7:**
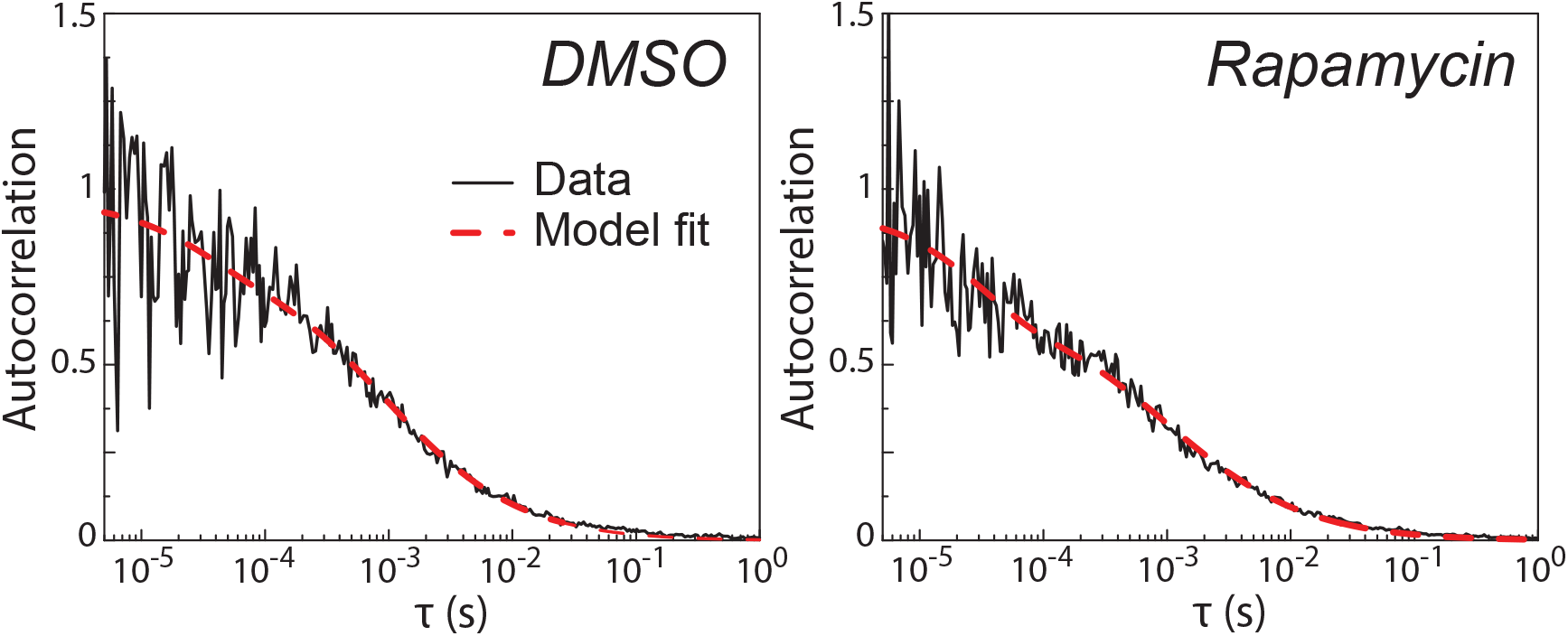
Fluorescence correlation spectroscopy indicates that the diffusion coefficient of 2xGFP is not affected by inhibition of mTORC1. Related to figure 2. Autocorrelation data of 2xGFP cells treated with DMSO vehicle control (A) or rapamycin (B), together with the fit using a blinking and anomalous diffusion model.

**Figure S8.**
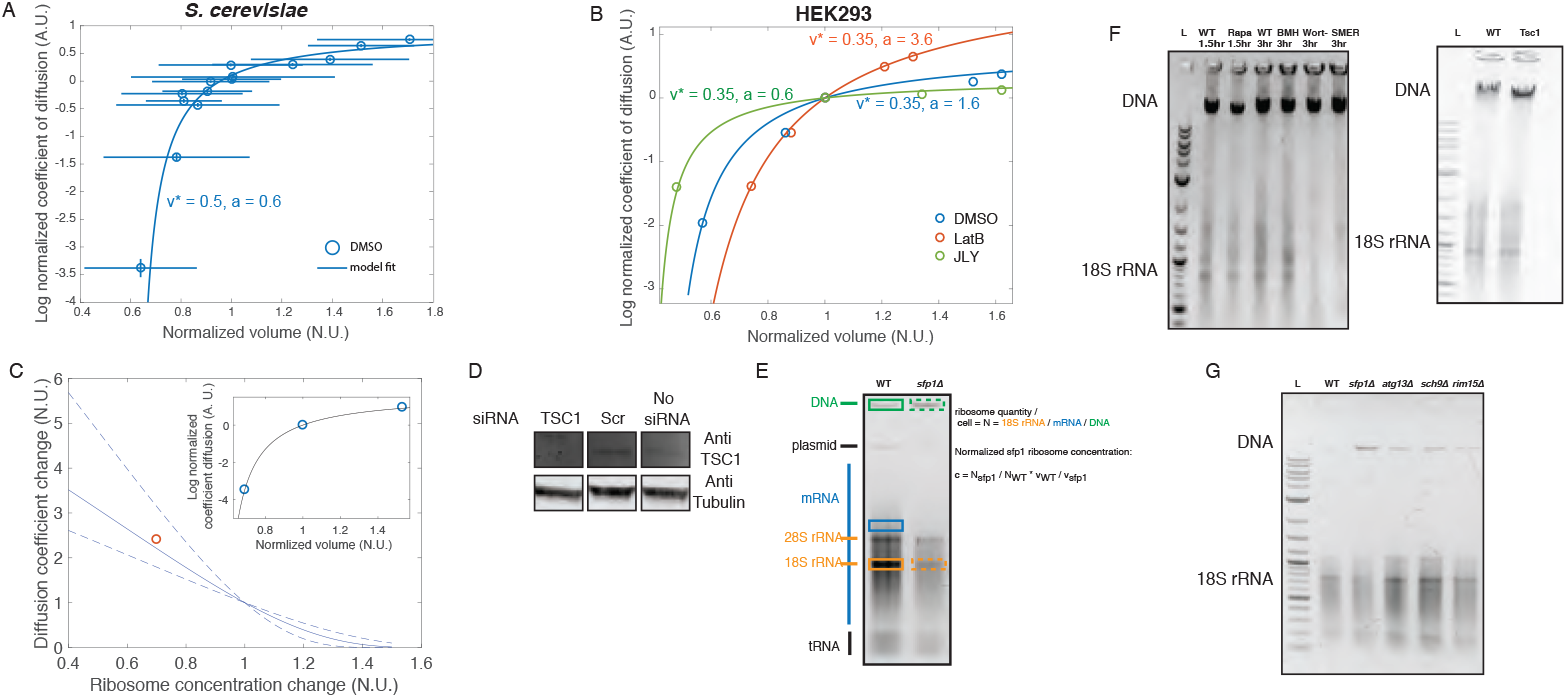
Validation of the Doolittle equation and determination of parameters using instantaneous volume change through osmotic stress, TSC western blot and 18S rRNA quantification, related to figures 3, 5 and 6. (**A**) After performing hyper- and hypo-osmotic shocks to perturb cell volume and then immediately assessed the diffusion coefficient for 40nm-GEMs, we fitted the model (equation (S10) on *S. cerevisiae* and find that it is in very good agreement with our data, suggesting that the Doolittle equation reasonably describes the dependence of diffusion coefficient on volume fraction of crowding agent (r^2^ = 0.85), and supplying parameters ζ ~ 0.6, *φ_0_/φ_m_* ~ 0.5. (**B**) We performed the same osmotic stress experiment to mammalian cells, and initially measured different parameters (ζ ~ 1.6, *φ_0_/φ_m_* ~ 0.35). Osmotic stress is known to strongly affect the actin cytoskeleton in mammalian cells, which was confirmed when we treated the cells with LatA at the same time we did the osmotic stress (ζ ~ 3.6, *φ_0_/φ_m_* ~ 0.35): the ζ interaction parameter of the GEMs with the environment increased. When the actin cytoskeleton was stabilized with JLY cocktail, we found that the 2 parameters of the model were closer to the yeast values: ζ ~ 0.6, is very similar to yeast, suggesting that the interactions of the GEMs with the microenvironment is the same, and *φ_0_/φ_m_* ~ 0.35 is lower, suggesting that mammalian cells are less crowded. (**C**) The non-osmotic volume and the ζ parameter for cells containing mRNP (inset) were calibrated in order to predict, for this type of particle, how the change in diffusion coefficient would be affected by a change in ribosome concentration, occurring through a rapamycin treatment. (**D**) TSC1 was targeted using Silencer Select siRNA (Thermo Fisher). Knockdown was validated by western blot using Hamartin/TSC1 with a tubulin control using standard techniques (see methods). (**E**) We extracted total nucleic acid by neutral phenol (see methods) and run the extract in a agarose gel, for the various chemical of genetic conditions explored. The gel is decomposed in the DNA band, that is used as a proxy for the amount of cells extracted, mRNA, rRNA and tRNA. To assess the relative amount of rRNA, as a proxy for ribosome amount, we normalized the band of rRNA to mRNA, and subsequently to DNA to get this quantity per cell. This number was extracted for each conditions, and normalized to the control: this gives us the relative change in ribosome number in HEK293 drug and siRNA treatments (**F**) and yeast mutants (**G**).

**Table S2: Change in basal diffusion and in epistasis with rapamycin for various mutations. Related to figure 3.**

Table S2 presents the nomenclature and a brief gene description of the various mutants explored in this study. The two numbers presented are the change in basal gene diffusion, measured as the ratio of 40nm-GEM diffusion in the mutant as compared to the wild type:

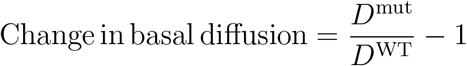

and the epistatic effect with rapamycin, measured as the ratio change of the diffusion under rapamycin for the mutant *v.* the wild type (equation formulated such that 0 indicates no epistasis and 1 indicates complete epistasis):

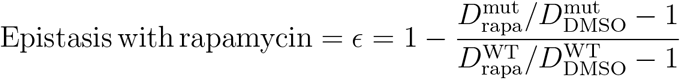

**Table S2:**
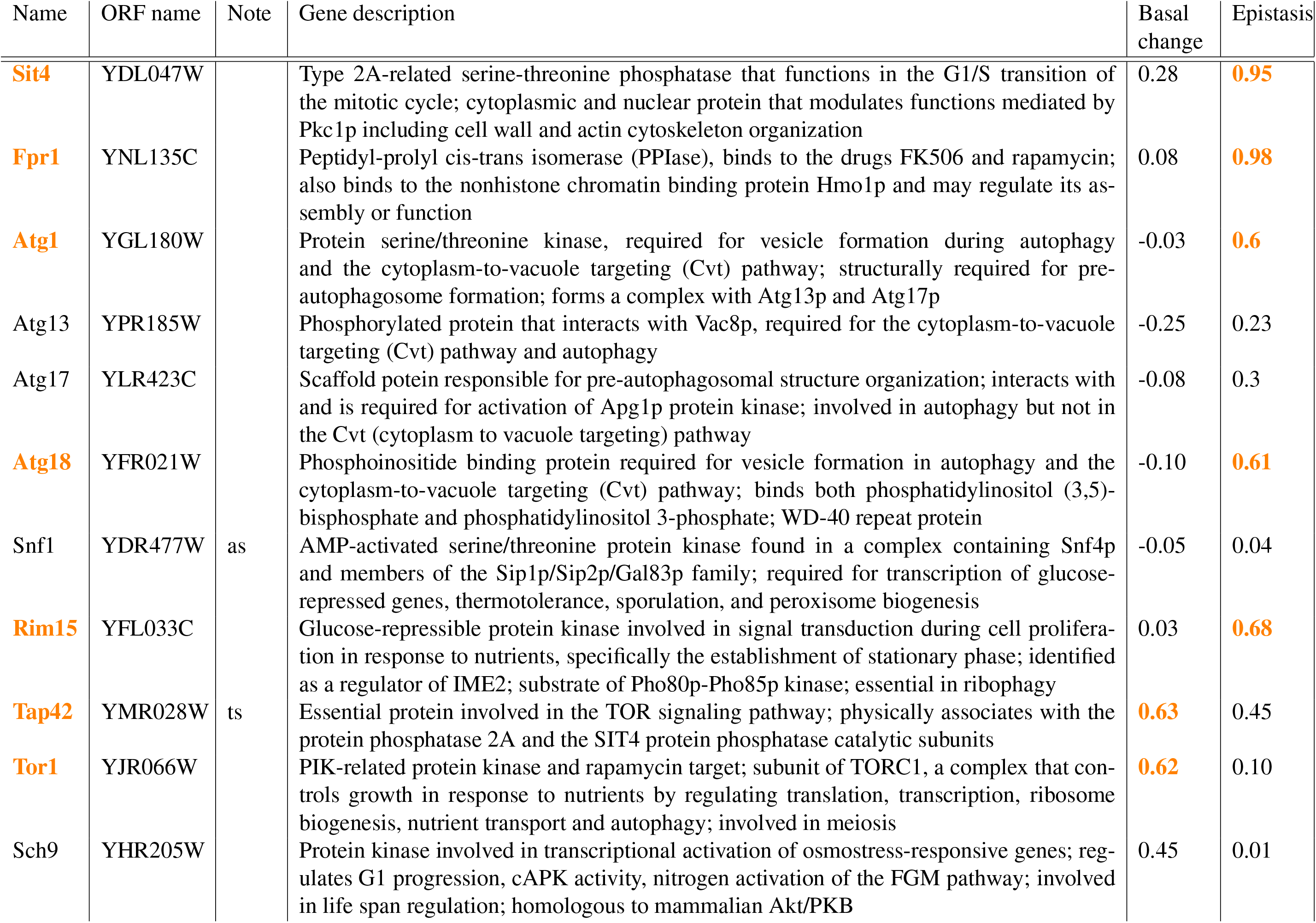
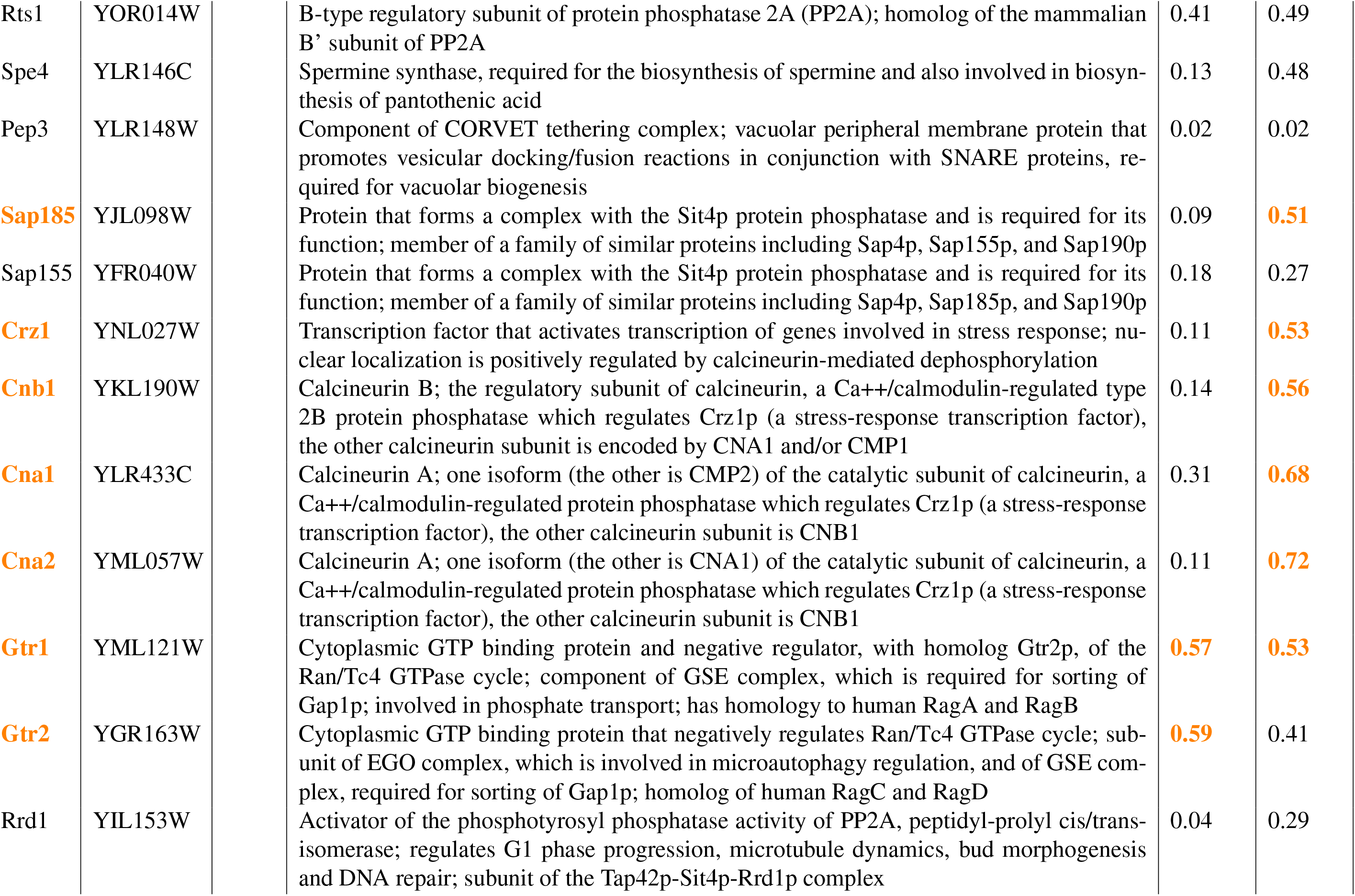
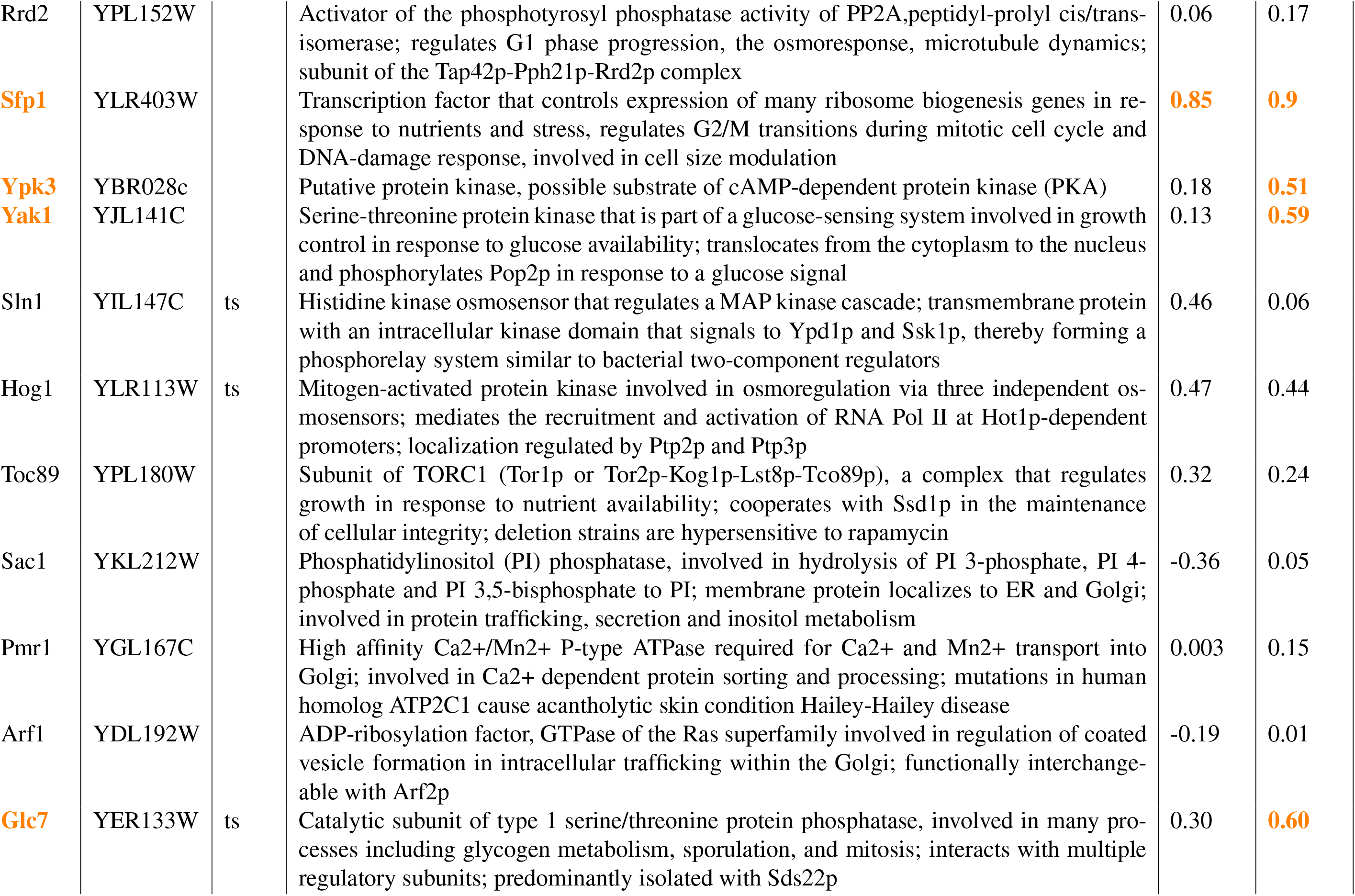
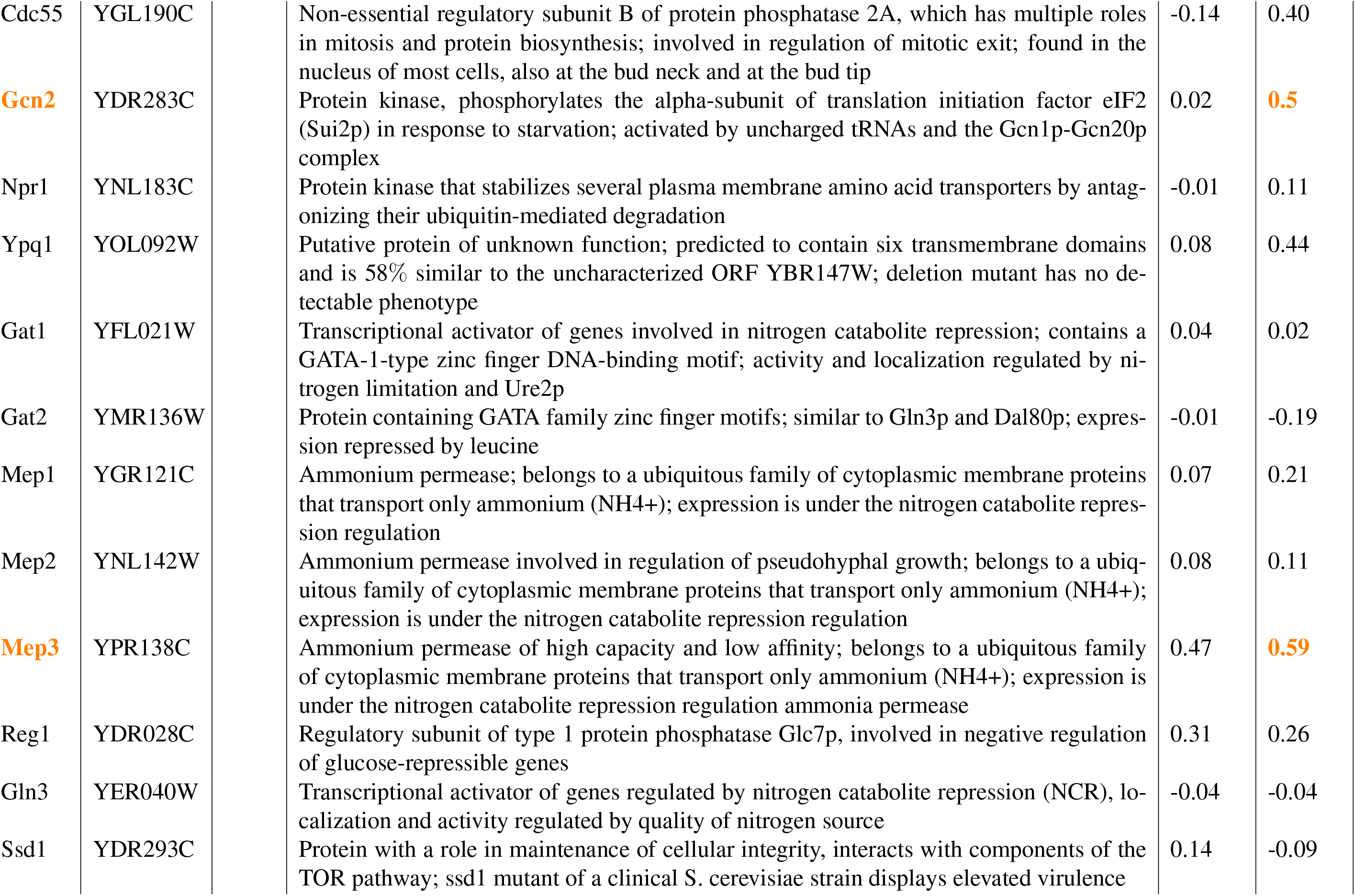
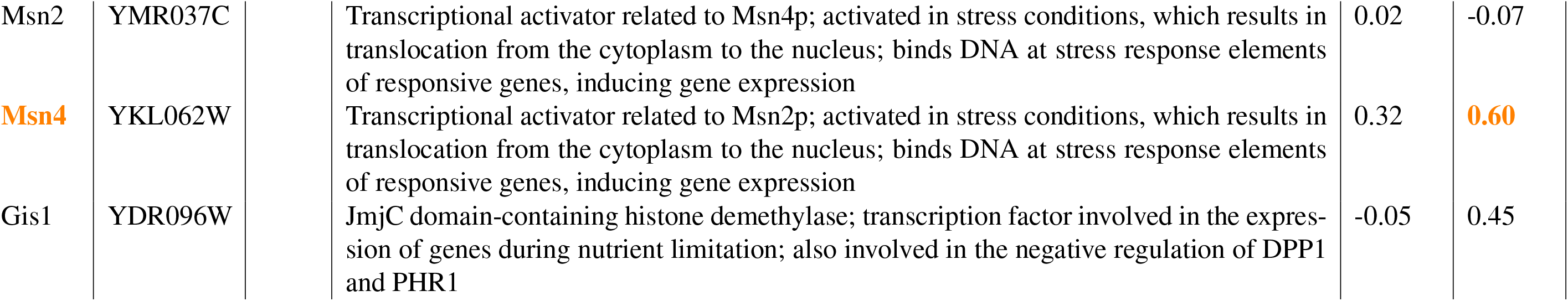
Mutants in the BY4741 background expressing pLH0497. Basal change in diffusion coefficient, and epistatic effect with rapamycin were measured. Significant differences, defined when the basal change, or the epistasis effect are higher than 0.5, are highlighted in orange, as well as the corresponding gene.

| Name | ORF name | Note | Gene description | Basal change | Epistasis |
| --- | --- | --- | --- | --- | --- |
| **Sit4** | YDL047W |  | Type 2A-related serine-threonine phosphatase that functions in the G1/S transition of the mitotic cycle; cytoplasmic and nuclear protein that modulates functions mediated by Pkc1p including cell wall and actin cytoskeleton organization | 0.28 | **0.95** |
| **Fpr1** | YNL135C |  | Peptidyl-prolyl cis-trans isomerase (PPIase), binds to the drugs FK506 and rapamycin; also binds to the nonhistone chromatin binding protein Hmo1p and may regulate its assembly or function | 0.08 | **0.98** |
| **Atg1** | YGL180W |  | Protein serine/threonine kinase, required for vesicle formation during autophagy and the cytoplasm-to-vacuole targeting (Cvt) pathway; structurally required for pre-autophagosome formation; forms a complex with Atg13p and Atg17p | -0.03 | **0.6** |
| Atg13 | YPR185W |  | Phosphorylated protein that interacts with Vac8p, required for the cytoplasm-to-vacuole targeting (Cvt) pathway and autophagy | -0.25 | 0.23 |
| Atg17 | YLR423C |  | Scaffold protein responsible for pre-autophagosomal structure organization; interacts with and is required for activation of Apg1p protein kinase; involved in autophagy but not in the Cvt (cytoplasm to vacuole targeting) pathway | -0.08 | 0.3 |
| **Atg18** | YFR021W |  | Phosphoinositide binding protein required for vesicle formation in autophagy and the cytoplasm-to-vacuole targeting (Cvt) pathway; binds both phosphatidylinositol (3,5)-bisphosphate and phosphatidylinositol 3-phosphate; WD-40 repeat protein | -0.10 | **0.61** |
| Snf1 | YDR477W | as | AMP-activated serine/threonine protein kinase found in a complex containing Snf4p and members of the Sip1p/Sip2p/Gal83p family; required for transcription of glucose-repressed genes, thermotolerance, sporulation, and peroxisome biogenesis | -0.05 | 0.04 |
| **Rim15** | YFL033C |  | Glucose-repressible protein kinase involved in signal transduction during cell proliferation in response to nutrients, specifically the establishment of stationary phase; identified as a regulator of IME2; substrate of Pho80p-Pho85p kinase; essential in ribophagy | 0.03 | **0.68** |
| **Tap42** | YMR028W | ts | Essential protein involved in the TOR signaling pathway; physically associates with the protein phosphatase 2A and the SIT4 protein phosphatase catalytic subunits | **0.63** | 0.45 |
| **Tor1** | YJR066W |  | PIK-related protein kinase and rapamycin target; subunit of TORC1, a complex that controls growth in response to nutrients by regulating translation, transcription, ribosome biogenesis, nutrient transport and autophagy; involved in meiosis | **0.62** | 0.10 |
| Sch9 | YHR205W |  | Protein kinase involved in transcriptional activation of osmostress-responsive genes; regulates G1 progression, cAPK activity, nitrogen activation of the FGM pathway; involved in life span regulation; homologous to mammalian Akt/PKB | 0.45 | 0.01 |

|  |  |  |  |  |
| --- | --- | --- | --- | --- |
| Rts1 | YOR014W | B-type regulatory subunit of protein phosphatase 2A (PP2A); homolog of the mammalian B' subunit of PP2A | 0.41 | 0.49 |
| Spe4 | YLR146C | Spermine synthase, required for the biosynthesis of spermine and also involved in biosynthesis of pantothenic acid | 0.13 | 0.48 |
| Pep3 | YLR148W | Component of CORVET tethering complex; vacuolar peripheral membrane protein that promotes vesicular docking/fusion reactions in conjunction with SNARE proteins, required for vacuolar biogenesis | 0.02 | 0.02 |
| **Sap185** | YJL098W | Protein that forms a complex with the Sit4p protein phosphatase and is required for its function; member of a family of similar proteins including Sap4p, Sap155p, and Sap190p | 0.09 | **0.51** |
| Sap155 | YFR040W | Protein that forms a complex with the Sit4p protein phosphatase and is required for its function; member of a family of similar proteins including Sap4p, Sap185p, and Sap190p | 0.18 | 0.27 |
| **Crz1** | YNL027W | Transcription factor that activates transcription of genes involved in stress response; nuclear localization is positively regulated by calcineurin-mediated dephosphorylation | 0.11 | **0.53** |
| **Cnb1** | YKL190W | Calcineurin B; the regulatory subunit of calcineurin, a Ca<sup>++</sup>/calmodulin-regulated type 2B protein phosphatase which regulates Crz1p (a stress-response transcription factor), the other calcineurin subunit is encoded by CNA1 and/or CMP1 | 0.14 | **0.56** |
| **Cna1** | YLR433C | Calcineurin A; one isoform (the other is CMP2) of the catalytic subunit of calcineurin, a Ca<sup>++</sup>/calmodulin-regulated protein phosphatase which regulates Crz1p (a stress-response transcription factor), the other calcineurin subunit is CNB1 | 0.31 | **0.68** |
| **Cna2** | YML057W | Calcineurin A; one isoform (the other is CNA1) of the catalytic subunit of calcineurin, a Ca<sup>++</sup>/calmodulin-regulated protein phosphatase which regulates Crz1p (a stress-response transcription factor), the other calcineurin subunit is CNB1 | 0.11 | **0.72** |
| **Gtr1** | YML121W | Cytoplasmic GTP binding protein and negative regulator, with homolog Gtr2p, of the Ran/Tc4 GTPase cycle; component of GSE complex, which is required for sorting of Gap1p; involved in phosphate transport; has homology to human RagA and RagB | **0.57** | **0.53** |
| **Gtr2** | YGR163W | Cytoplasmic GTP binding protein that negatively regulates Ran/Tc4 GTPase cycle; subunit of EGO complex, which is involved in microautophagy regulation, and of GSE complex, required for sorting of Gap1p; homolog of human RagC and RagD | **0.59** | 0.41 |
| Rrd1 | YIL153W | Activator of the phosphotyrosyl phosphatase activity of PP2A, peptidyl-prolyl cis/trans-isomerase; regulates G1 phase progression, microtubule dynamics, bud morphogenesis and DNA repair; subunit of the Tap42p-Sit4p-Rrd1p complex | 0.04 | 0.29 |
| Rrd2 | YPL152W |  | Activator of the phosphotyrosyl phosphatase activity of PP2A, peptidyl-prolyl cis/trans-isomerase; regulates G1 phase progression, the osmoresponse, microtubule dynamics; subunit of the Tap42p-Pph21p-Rrd2p complex | 0.06 | 0.17 |
| **Sfp1** | YLR403W |  | Transcription factor that controls expression of many ribosome biogenesis genes in response to nutrients and stress, regulates G2/M transitions during mitotic cell cycle and DNA-damage response, involved in cell size modulation | **0.85** | **0.9** |
| **Ypk3** | YBR028c |  | Putative protein kinase, possible substrate of cAMP-dependent protein kinase (PKA) | 0.18 | **0.51** |
| **Yak1** | YJL141C |  | Serine-threonine protein kinase that is part of a glucose-sensing system involved in growth control in response to glucose availability; translocates from the cytoplasm to the nucleus and phosphorylates Pop2p in response to a glucose signal | 0.13 | **0.59** |
| Sln1 | YIL147C | ts | Histidine kinase osmosensor that regulates a MAP kinase cascade; transmembrane protein with an intracellular kinase domain that signals to Ypd1p and Ssk1p, thereby forming a phosphorelay system similar to bacterial two-component regulators | 0.46 | 0.06 |
| Hog1 | YLR113W | ts | Mitogen-activated protein kinase involved in osmoregulation via three independent osmosensors; mediates the recruitment and activation of RNA Pol II at Hot1p-dependent promoters; localization regulated by Ptp2p and Ptp3p | 0.47 | 0.44 |
| Toc89 | YPL180W |  | Subunit of TORC1 (Tor1p or Tor2p-Kog1p-Lst8p-Tco89p), a complex that regulates growth in response to nutrient availability; cooperates with Ssd1p in the maintenance of cellular integrity; deletion strains are hypersensitive to rapamycin | 0.32 | 0.24 |
| Sac1 | YKL212W |  | Phosphatidylinositol (PI) phosphatase, involved in hydrolysis of PI 3-phosphate, PI 4-phosphate and PI 3,5-bisphosphate to PI; membrane protein localizes to ER and Golgi; involved in protein trafficking, secretion and inositol metabolism | -0.36 | 0.05 |
| Pmr1 | YGL167C |  | High affinity Ca<sup>2+</sup>/Mn<sup>2+</sup> P-type ATPase required for Ca<sup>2+</sup> and Mn<sup>2+</sup> transport into Golgi; involved in Ca<sup>2+</sup> dependent protein sorting and processing; mutations in human homolog ATP2C1 cause acantholytic skin condition Hailey-Hailey disease | 0.003 | 0.15 |
| Arf1 | YDL192W |  | ADP-ribosylation factor, GTPase of the Ras superfamily involved in regulation of coated vesicle formation in intracellular trafficking within the Golgi; functionally interchangeable with Arf2p | -0.19 | 0.01 |
| **Glc7** | YER133W | ts | Catalytic subunit of type 1 serine/threonine protein phosphatase, involved in many processes including glycogen metabolism, sporulation, and mitosis; interacts with multiple regulatory subunits; predominantly isolated with Sds22p | 0.30 | **0.60** |

Table S2: Mutants in the BY4741 background expressing pLH0497.
|  |  |  |  |  |
| --- | --- | --- | --- | --- |
| Cdc55 | YGL190C | Non-essential regulatory subunit B of protein phosphatase 2A, which has multiple roles in mitosis and protein biosynthesis; involved in regulation of mitotic exit; found in the nucleus of most cells, also at the bud neck and at the bud tip | -0.14 | 0.40 |
| **Gcn2** | YDR283C | Protein kinase, phosphorylates the alpha-subunit of translation initiation factor eIF2 (Sui2p) in response to starvation; activated by uncharged tRNAs and the Gcn1p-Gcn20p complex | 0.02 | **0.5** |
| Npr1 | YNL183C | Protein kinase that stabilizes several plasma membrane amino acid transporters by antagonizing their ubiquitin-mediated degradation | -0.01 | 0.11 |
| Ypq1 | YOL092W | Putative protein of unknown function; predicted to contain six transmembrane domains and is 58% similar to the uncharacterized ORF YBR147W; deletion mutant has no detectable phenotype | 0.08 | 0.44 |
| Gat1 | YFL021W | Transcriptional activator of genes involved in nitrogen catabolite repression; contains a GATA-1-type zinc finger DNA-binding motif; activity and localization regulated by nitrogen limitation and Ure2p | 0.04 | 0.02 |
| Gat2 | YMR136W | Protein containing GATA family zinc finger motifs; similar to Gln3p and Dal80p; expression repressed by leucine | -0.01 | -0.19 |
| Mep1 | YGR121C | Ammonium permease; belongs to a ubiquitous family of cytoplasmic membrane proteins that transport only ammonium (NH<sub>4</sub><sup>+</sup>); expression is under the nitrogen catabolite repression regulation | 0.07 | 0.21 |
| Mep2 | YNL142W | Ammonium permease involved in regulation of pseudohyphal growth; belongs to a ubiquitous family of cytoplasmic membrane proteins that transport only ammonium (NH<sub>4</sub><sup>+</sup>); expression is under the nitrogen catabolite repression regulation | 0.08 | 0.11 |
| **Mep3** | YPR138C | Ammonium permease of high capacity and low affinity; belongs to a ubiquitous family of cytoplasmic membrane proteins that transport only ammonium (NH<sub>4</sub><sup>+</sup>); expression is under the nitrogen catabolite repression regulation ammonia permease | 0.47 | **0.59** |
| Reg1 | YDR028C | Regulatory subunit of type 1 protein phosphatase Glc7p, involved in negative regulation of glucose-repressible genes | 0.31 | 0.26 |
| Gln3 | YER040W | Transcriptional activator of genes regulated by nitrogen catabolite repression (NCR), localization and activity regulated by quality of nitrogen source | -0.04 | -0.04 |
| Ssd1 | YDR293C | Protein with a role in maintenance of cellular integrity, interacts with components of the TOR pathway; ssd1 mutant of a clinical *S. cerevisiae* strain displays elevated virulence | 0.14 | -0.09 |
| Msn2 | YMR037C | Transcriptional activator related to Msn4p; activated in stress conditions, which results in translocation from the cytoplasm to the nucleus; binds DNA at stress response elements of responsive genes, inducing gene expression | 0.02 | -0.07 |
| **Msn4** | YKL062W | Transcriptional activator related to Msn2p; activated in stress conditions, which results in translocation from the cytoplasm to the nucleus; binds DNA at stress response elements of responsive genes, inducing gene expression | 0.32 | **0.60** |
| Gis1 | YDR096W | JmjC domain-containing histone demethylase; transcription factor involved in the expression of genes during nutrient limitation; also involved in the negative regulation of DPP1 and PHR1 | -0.05 | 0.45 |

